# Regulation of Kv1.2 redox-sensitive gating by the transmembrane lectin LMAN2

**DOI:** 10.1101/2024.07.09.602766

**Authors:** Shawn M. Lamothe, Damayantee Das, Anson A. Wong, Yubin Hao, Aislinn D. Maguire, Bradley J. Kerr, Victoria A. Baronas, Harley T. Kurata

**Affiliations:** Dept. of Pharmacology, Alberta Diabetes Institute, University of Alberta, 9-70 Medical Sciences Building, Edmonton, AB, T6G 2H7, Canada; Neuroscience and Mental Health Institute, University of Alberta, Edmonton, AB, Canada; Dept. of Anesthesiology and Pain Medicine, University of Alberta, Edmonton, AB, Canada; Dept. of Surgery, University of British Columbia, Vancouver, BC, Canada

**Author notes:** Corresponding author: Harley T. Kurata, Current address: Dept. of Pharmacology, Alberta Diabetes Institute, University of Alberta, 9-70 Medical Sciences Building, Edmonton, AB, T6G 2H7, Canada.

## Abstract

Kv1.2 potassium channels influence excitability and action potential propagation in the nervous system. Unlike closely-related Kv1 channels, Kv1.2 exhibits highly variable voltage-dependence of gating, attributed to regulation by unidentified extrinsic factors. Variable Kv1.2 gating is strongly influenced by the extracellular redox potential, and we demonstrate that Kv1.2 currents in dorsal root ganglion sensory neurons exhibit similar variability and redox sensitivity as observed when the channel is heterologously expressed in cell lines. We used a functional screening approach to test the effects of candidate regulatory proteins on Kv1.2 gating, using patch clamp electrophysiology. Among 52 candidate genes tested, we observed that co-expression with the transmembrane lectin LMAN2 led to a pronounced gating shift of Kv1.2 activation to depolarized voltages in CHO and L(tk-) cell lines, caused by deceleration of activation kinetics. Overexpression of LMAN2 promoted a slow gating mode of Kv1.2 that mimics the functional outcomes of extracellular reducing conditions, and enhanced sensitivity to extracellular reducing agents. In contrast, shRNA-mediated knockdown of endogenous LMAN2 in cell lines reduced Kv1.2 redox sensitivity and gating variability. Kv1.2 sensitivity to LMAN2 is abolished by mutation of neighboring residues F251 and T252 in the intracellular S2-S3 linker, and these also abolish redox-dependent modulation of Kv1.2, suggesting that LMAN2 is an important contributor to the mechanism of redox sensitivity. In conclusion, we identified LMAN2 as a candidate regulatory protein that influences redox-dependent modulation of Kv1.2, and clarified the structural elements of the channel that are required for sensitivity.

## INTRODUCTION

Voltage gated potassium (Kv) channels have a pivotal role in the regulation of threshold excitability and action potential morphology in excitable cells (Shen *et al*., 2004; Goldberg *et al*., 2008). Voltage-gated channels may experience varied patterns of stimulation (eg. tonic vs. phasic firing) where frequency dependent responses shape cellular behavior. Kv1.2 is unique among most Kv channels because it exhibits ‘use-dependent activation’, where trains of repetitive depolarizations influence the kinetics and voltage-dependence of activation, and this may alter threshold and burst firing properties of action potentials (Rezazadeh *et al*., 2007; Baronas *et al*., 2016). While the physiological role for Kv1.2 use-dependent activation has not been determined, altered function of Kv1.2 in the CNS causes a variety of neurological phenotypes including ataxia, epilepsy and encephalopathy (Xie *et al*., 2010; Pena and Coimbra, 2015; Corbett *et al*., 2016; Masnada *et al*., 2017; Sachdev *et al*., 2017). In addition, downregulation of Kv1.2 currents contributes to hyperexcitability in dorsal root ganglion (DRG) neurons and the emergence of neuropathic pain (Ishikawa *et al*., 1999; Yang *et al*., 2004). Thus, characterizing the mechanisms regulating Kv1.2 channels is important for understanding physiological electrical signaling and its dysregulation in neurological diseases.

Kv1.2 use-dependent activation arises from channels switching between a slow ‘inhibited’ gating mode and a faster ‘potentiated’ gating mode. Previous studies have demonstrated that this process is sensitive to extracellular redox potential, and it is likely that this mode switching is mediated by Kv1.2 interactions with unknown extrinsic factors such as accessory proteins (Rezazadeh *et al*., 2007; Baronas *et al*., 2015, 2016). An important related observation is that Kv1.2 gating exhibits prominent cell-to-cell variability in heterologous cell lines and in primary neuronal cultures, indicating the channel itself does not contain all necessary elements for use-dependent gating, and that other endogenously expressed regulatory elements are involved (Baronas *et al*., 2015). Previously reported Kv1.2 regulators such as Kvβ subunits, C-terminal phosphorylation, σ-1 receptor, and calnexin, do no account for the pronounced gating variability of Kv1.2 (Rettig *et al*., 1994; Yang *et al*., 2007; Stirling *et al*., 2009; Kourrich *et al*., 2013). Recent progress from our group has uncovered additional extrinsic factors that regulate Kv1.2 gating. We reported that a neutral amino acid transporter, Slc7a5, causes a dramatic hyperpolarizing shift (∼-50 mV) of Kv1.2 voltage of half-activation (V_1/2_), while suppressing current magnitude of both Kv1.1 and Kv1.2 (Baronas *et al*., 2018; Lamothe and Kurata, 2020; Lamothe *et al*., 2020). In contrast, use-dependent activation is markedly enhanced together with a prominent *depolarizing* shift of the voltage of half-activation (V_1/2_) by small concentrations of extracellular reducing agents (e.g., DTT and TCEP, which cause a roughly 80 mV depolarizing shift, leading to a V_1/2_ of ∼60 mV). Importantly, this redox sensitivity does not involve native cysteine residues in Kv1.2, offering further evidence that additional redox sensitive proteins extrinsic to the channel are involved (Baronas *et al*., 2017). Thus, while evidence has accumulated describing prominent modulation of use- and voltage-dependent activation of Kv1.2, the molecular mechanisms attributing to this behavior and the underlying redox-sensitive modulation are still not understood.

In this study we explored the redox sensitivity of Kv currents encoded by Kv1.2-containing channels in primary DRG neurons. We also investigated potential mechanisms of Kv1.2 redox sensitivity by screening for Kv1.2 modulation by previously unrecognized candidate proteins. We report that the transmembrane lectin LMAN2 (VIP36), dramatically influences Kv1.2 redox-sensitive gating in multiple cell lines. Co-expression of LMAN2 with Kv1.2 in heterologous cell lines shifts the voltage-dependence of Kv1.2 activation to strong positive voltages, and enhances channel sensitivity to extracellular reducing conditions, thereby mimicking the gating effects of extracellular reducing agents. Alternatively, shRNA-mediated knockdown of LMAN2, diminishes Kv1.2 use-dependence and redox sensitivity. We localized an overlapping determinant of LMAN2 and redox sensitivity of Kv1.2 to the intracellular S2-S3 linker, a region previously shown to be crucial for Kv1.2 use-dependent activation (Rezazadeh *et al*., 2007; Baronas *et al*., 2016). We also established that unlike other members of the mammalian L-type transmembrane lectin family (LMAN1 and LMAN2L), LMAN2 shuttling to the cell-surface is an important determinant for Kv1.2 regulation. In summary, we have identified a previously unrecognized protein that strongly influences Kv1.2 redox sensitivity and promotes use-dependent activation.

## MATERIALS AND METHODS

### Cell Culture and Expression

Mouse L(tk-) fibroblast cells (ATCC CCL-1.3) (‘LM cells’) were used for patch clamp electrophysiology, shRNA stable cell line generation, Western blots, and qPCR. Chinese Hamster Ovary (CHO) cells were used for patch clamp electrophysiology. HEK-293 cells were used for Western blots. Cells were maintained in culture in a 5% CO_2_ incubator at 37°C in complete medium (DMEM high glucose (Sigma: D5796) + 10% FBS + 1% penicillin/streptomycin). For patch clamp electrophysiology and Western blotting, cells were split into 12-well plates to achieve 70% confluency, then transfected with cDNA for 24-48 hours using jetPRIME® transfection reagent (Polyplus) as per manufacturer’s instructions. Fluorescent proteins were either fused or co-transfected with cDNA of interest to identify transfected cells for electrophysiological recording. Unless otherwise indicated, cells were consistently transfected with 1:4 ratios of channel DNA to either N-terminally tagged EGFP constructs or EGFP to maintain a constant amount of DNA in the transfection mixture. 6-10 hours after transfection, cells were split onto glass coverslips for electrophysiological recordings from single cells the following day.

Electrophysiological recordings were done 24-48 hours after transfection. Western blot experiments were performed 48-72 hours after transfection.

### Plasmids and mutagenesis

All Kv1.2-encoding plasmids (including various Kv1.2 mutants and chimeras) were expressed using the pcDNA3.1(-) plasmid. The rat isoform of Kv1.2 (NM_012970.4) was used throughout the study. Candidate regulatory proteins were obtained from the DNASU plasmid repository, and subcloned for expression in the pEGFP-C1 plasmid, so all were expressed with N-terminal EGFP tags unless explicitly stated otherwise (i.e. Fig. 8). The Kv1.2 [T252R] point mutant has been previously described (Baronas *et al*., 2017). Mutagenesis of Kv1.2 and LMAN2 to generate Kv1.2 [F251S] and LMAN2 [KKAA], respectively, was carried out using two-step overlapping PCR in the mutated region, followed by verification with Sanger sequencing. The following primers were used to generate these constructs:

Kv1.2[F251S]

Forward, 5’-GCTGGCTTCAGCACCAACATC-3’

Reverse, 5’ CGACCGAAGTCGTGGTTGTAG-3’

LMAN2 [KKAA]

Forward, 5’-GGGGCTAGCATGGTGAGCAAGGGCGAGGAG-3’

Reverse, 5’-GGGGAATTCTCAAGCAGCTTTCTTGTTCCGCTCCTGCCGCTTC-3’

Kv1.2/Kv1.5 chimeras replacing the S1 segment, S1-S2 Linker, S2 segment and S2-S3L of Kv1.2 with that of Kv1.5, along with the respective primers used to generate the chimeras, were previously described (Lamothe *et al*., 2020). These chimeras correspond to substitution of Kv1.2 sequence with Kv1.5 between amino acid positions (Kv1.2 numbering): 153-187 (Kv1.2 (1.5 S1)); 187-217 (Kv1.2 (1.5 S1-S2L)); 217-242 (Kv1.2 (1.5 S2)); 242-259 (Kv1.2 (1.5 S2-S3L)).

The Kv1.2 (1.5 S1-S3) chimera was constructed from a previously described chimera comprising the Kv1.2 N-terminus followed by the VSD, pore, and C-terminus of Kv1.5 (Lamothe *et al*., 2020). PCR of the N-terminus until the intracellular side of S3 of this chimera was used to generate Fragment 1. PCR of the S3 segment to the C-terminus of Kv1.2 was used to generate Fragment 2. Overlapping PCR of Fragments 1 and 2 generated a chimeric channel comprising Kv1.2, with Kv1.5 sequence substituted from residues 153-259 (Kv1.2 numbering. This was cloned into pcDNA3.1(-) with NheI and HindIII restriction enzymes. The following primers were used to generate this construct:

Fragment 1:

Forward, 5’-GGGGCTAGCATGACAGTGGCTACCGGAG-3’ (Kv1.2 NheI 5’)

Reverse, 5’-GAAGTAGGGGAAGATGGCCAC-3’

Fragment 2:

Forward, 5’-ATGAACATCATTGACATTGTGGCTATCATCCCTTAC-3’

Reverse, 5’-GGGAAGCTTTCAGACATCAGTTAAC-3’ (Kv1.2 HindIII 3’)

LMAN2 and other lectins (LMAN1 and LMAN2L) were expressed using the pcDNA3.1(-) vector. EGFP-tagged versions were generated by PCR amplification of the lectin gene and EGFP with compatible restriction sites followed by ligation into the multiple coding sequence of pcDNA3.1(-).

### Dorsal Root Ganglion (DRG) neuron isolation

Mice were euthanized by intraperitoneal pentobarbital overdose (Euthansol, 0.1 ml of 340 mg/ml) followed by intracardiac flush with 0.9% saline. Lumbar DRGs (L3-L6) were extracted from euthanized animals promptly. Immediately after extraction, mice lumbar DRGs (L3-L6) were immersed in ice-cold dissection solution (118 mM NaCl, 2.5 mM KCl, 1.3 μM MgSO_4_, 1.2 mM NaH_2_PO_4_, 5 mM MgCl_2_6H_2_O, 25 mM D-glucose, 26 mM NaHCO_3_, and 1.5 mM CaCl_2_). Shortly thereafter, DRGs were digested with 0.5-mg/ml trypsin (Sigma, catalogue number T-9201), 1 mg/ml collagenase Type IV (Cedarlane, catalogue number LS004186), and 0.1 mg/ml deoxyribonuclease I (Sigma, catalogue number D-5025) dissolved in DMEM supplemented with GlutaMax (Invitrogen, catalogue number 10569044) for 40 min in a shaking water bath set at 35°C. Dissociated cells were plated onto 35 × 10 mm dishes (VWR, catalogue number CA25382331) that were pretreated with 3 μg/ml poly-DL-ornithine (Sigma, catalogue number P-8638) dissolved in HPLC water (Sigma) and 2-μg/ml laminin (Sigma, catalogue number L-2020) dissolved in HBSS (138 mM NaCl, 5.33 mM KCl, 0.44 mM KH_2_PO_4_, 0.5 mM MgCl_2_ 6H_2_O, 0.41 mM MgSO_4_ 7H_2_O, 4 mM NaHCO_3_, 0.3 mM Na_2_HPO_4_, 5.6 mM D-glucose, and 1.26 mM CaCl_2_). Each 35 × 10 mm dish was immersed in 2 ml of culture medium (20 ml total), which contained 18-ml DMEM+GlutaMax (Invitrogen, catalogue number 10569044), 2-ml heat-inactivated horse serum (Sigma, catalogue number H-1138), 200 μl antibiotic-antimycotic 100× (Invitrogen, catalogue number 15240-096), and 20 μl antimitotic [cytosine β-D-arabinofuranoside (Ara-C), uridine, 5-fluoro-2’-deoxyuridine all at 10 μM (Sigma, catalogue numbers C1768, U3003, and F0503)]. Finally, cells were incubated at 37.0°C, 95% air – 5% CO_2_ for 24 h. All animal experiments were performed with approval from the University of Alberta Health Sciences Animal Care and Use Committee (protocol AUP0000274).

### Electrophysiology

Patch pipettes were manufactured from soda lime capillary glass (Fisher), using a Sutter P-97 pipette puller (Sutter Instrument). When filled with standard recording solutions, pipettes had a tip resistance of 1-3 MΩ. Recordings were filtered at 5 kHz, sampled at 10 kHz, with manual capacitance compensation and 80% series resistance compensation. Recordings were stored directly onto a computer hard drive using Clampex 10 software (Molecular Devices). For recording of Kv currents in heterologous cells, external (bath) recording solution had the following composition (in mM): 135 NaCl, 5 KCl, 1 CaCl_2_, 1 MgCl_2_, 10 HEPES and adjusted to pH 7.4 with NaOH. The internal (pipette) recording solution had the following composition (in mM): 135 KCl, 5 K-EGTA, 10 HEPES and adjusted to pH 7.2 with KOH. For isolation of Kv currents in DRG neurons, external (bath) recording solution had the following composition (in mM): 137 NMDG, 5.4 KCl, 1 MgCl2, 0.02 Ca^2+^, 10 D-Glucose, 10 HEPES (pH 7.3 with HCl). The internal (pipette) recording solution had the following composition (in mM): 135 KCl, 5 K-EGTA, 10 HEPES (pH 7.2 with KOH). Chemicals for electrophysiological solutions were purchased from Sigma-Aldrich or Fisher. Tityustoxin-Kα (TsTX) was purchased from alomone labs (cat #: STT-360) (dissolved in ddH_2_O, stored at 50 μM concentration) then diluted in extracellular recording solution to 100 nM on the day of experiments. Dithiothreitol (DTT) was obtained from Fisher Scientific (BP172-25), prepared as a 200 mM stock solution in ddH_2_O and diluted in recording solutions to appropriate concentrations on the day of experiments. Tris-(2-Carboxyethyl)phosphine hydrochloride (TCEP-HCL) was obtained from Thermo Fisher Scientific (Product #20490), prepared as a 50 mM stock in ddH_2_O solution and adjusted to pH 7.2 with NaOH. The stock solution was diluted in external recording solution to the desired working concentration on the day of experiments.

For data analysis, use-dependent activation (UDA) of channels during repetitive trains of depolarization was calculated as %UDA = (1 - 1^st^ pulse magnitude/peak pulse magnitude) * 100. Conductance-voltage relationships were determined by normalizing the tail current magnitude (at -30 mV) and fitting with a Boltzmann equation of the form G = 1/(1+e^-(V-V_1/2_)/*k*), where G is macroscopic conductance, V is the voltage applied, V_1/2_ is the half-maximal activation voltage, and *k* is a slope factor reflecting the voltage range over which an e-fold change in open probability (Po) is observed. The sum of two Boltzmanns of the form G = a/(1+e^-(V-V_1/2,_a)/*k,a*) + (1-a)/(1+e^-(V-V_1/2,_b)/*k,b*) was occasionally required due to conductance-voltage relationships with both slow and fast gating components. Rise time (20-80% of peak) was used to assess kinetics of channel activation, using fitting functions in Clampfit. Box plots depict the median, 25^th^ and 75^th^ percentile (box), and 10^th^ and 90^th^ percentile (whiskers).

### Action potential recordings

For measurement of action potentials from mouse DRG neurons, recordings were generated in current-clamp mode using an Axopatch 200B amplifier (Molecular Devices). Whole-cell configuration was obtained in voltage-clamp mode before manually switching to current-clamp recording mode. Electrodes had a tip resistance of 3-6 MΩ when filled with pipette solution. For action potential recordings from DRG neurons, external (bath) recording solution had the following composition (in mM): 135 NaCl, 5 KCl, 1 CaCl_2_, 1 MgCl_2_, 10 HEPES and adjusted to pH 7.4 with NaOH. The internal (pipette) recording solution had the following composition (in mM): 135 KCl, 4 Na_2_-ATP, 5 K-EGTA, 10 HEPES and adjusted to pH 7.2 with KOH. The frequency of action potentials was analyzed using the event detection, Threshold search, feature of clampfit 10.7. The frequency of action potentials was calculated by the number of spikes over the time exposed to a particular condition (Ambient, TCEP or TsTX). The baseline was set at the membrane potential at the beginning of the recording. The threshold level (Trigger) for detection of an action potential was set at 0 mV.

### LMAN2 gene knockdown plasmid design

shRNAs were generated targeting mouse LMAN2 for knockdown in L(tk-) cells (RefSeq: NM_025828.3 Mus musculus lectin, mannose-binding 2), with the LMAN2 mouse 29mer shRNA constructs (Origene, cat #: TL516848) in a pGFP-C-shLenti vector according to the manufacturer’s instructions. Scrambled (control) shRNA oligos (Biosettia) were cloned into the pLV-RNAi vector system (Biosettia, San Diego, USA. Cat #: sort-B21) according to the manufacturer’s instructions, as described previously (Lamothe *et al*., 2020; Satou *et al*., 2020). The LMAN2-targeted shRNA sequences tested were:

TL516848A: ATCACCGACGGCAACAGCGAACACCTCAA ;

TL516848B: GCCTGTGTTTGGAAGCAAAGACAACTTCC ;

TL516848C: AGGAGAGCATCGACTGGACCAAGATTGAG ;

TL516848D: TGGACGATGGAGTGAGTTGGCAGGCTGCA ;

Scrambled (control): GCAGTTATCTGGAAGATCAGGTTGGATCCAACCTGATCTTCCAGATAACTGC

Evaluation of stable cell lines for effective LMAN2 knockdown led to selection of the TL516848B shRNA-expressing cell line for experimental studies.

### Kv1.2 stable cell line plasmid design

Kv1.2_1D4 LM fibroblast stable cell lines were generated by cloning the rat isoform of Kv1.2 (NM_012970.4) with a C-terminal 1D4 tag into the pLV-expression vector

(pLV-EF1ɑ-MCS-IRES-RFP-Puro, Biosettia, Cat #: cDNA-pLV09) with BamHI and NheI restriction enzymes,T4 DNA ligase and the following primers:

Forward, 5’-CCCGGATCCGCCACCATGACAGTGGCTACCGGAGAC-3’

Reverse, 5’-CCCGCTAGCTCACGCCGGCGCCACCTGG-3’

LMAN2 and scrambled shRNAs and Kv1.2_1D4 constructs were confirmed by Sanger sequencing (Applied Genomics Core, University of Alberta).

### Lentiviral preparation and delivery

shRNA vectors or Kv1.2_1D4 expression vectors and packaging vectors (pLV-PACK packaging mix, Biosettia, Cat #: pLV-PACK-500) were co-transfected into HEK293 cells with Lipofectamine2000 (Thermo Fisher Scientific) in 10 cm culture plates, and the cell culture supernatant was harvested after 48 hours. LM fibroblasts cells were seeded at approximately 30% confluence into a 6-well (30 mm) plate containing 2 ml of complete medium (DMEM with 10%FBS) 12-24 hours before viral transduction. Afterwards, the complete medium was removed and medium containing 8 μg/ml polybrene and 500 μl of HEK293 cell culture supernatant containing the virus was added to the LM cells. The virus-containing medium was replaced with fresh complete medium 24-hour after incubation. After further incubation for 48 hours, 2.5 μg/ml puromycin was added and maintained in cell culture for selection of successfully transduced cells. Effectiveness of knockdown or Kv1.2 overexpression was assessed using qPCR and Western blot approaches.

### Biotinylation/Isolation of cell-surface proteins

A cell surface protein isolation kit (Pierce) was used to determine the cell surface localization of LMAN2, LMAN1, LMAN2L and LMAN2[KKAA]. In brief, cells were grown to 60-80% confluence in 10 cm culture plates. Cells were transfected with 2 μg of the appropriate plasmids, 48 hours before cell-surface protein isolation. Membrane proteins were labeled with a membrane-impermeant thiol-cleavable amine-reactive biotinylating reagent, Sulfo-NHS-SS biotin, for 10 minutes at room-temperature then washed with ice-cold TBS. Treated cells were collected in 10 ml of TBS then cell pellets were lysed with a lysis buffer containing 1% protease inhibitor mixture. After centrifugation for 5 min at 15,000 x *g* at 4°C, biotin-labeled proteins were isolated using NeutrAvidin™ agarose columns and eluted with a buffer containing 1 mM DTT. The concentrations of isolated cell-surface and total proteins were determined and analyzed via Western blot analysis. As a loading control, Na^+^/K^+^-ATPase (ATP1A1) expression was detected using rabbit anti-Na^+^/K^+^-ATPase α (ATP1A1) primary antibody (cell signaling: #3010) and anti-rabbit HRP-conjugated secondary antibody (Abcam: ab6721). Total protein was detected, along with β-actin as the loading control (anti-β-actin primary antibody (Genetex: GT5512), anti-mouse HRP-conjugated secondary antibody (Applied Biological Materials: SH023)).

### Western blot

Detection of LMAN2, LMAN1 and LMAN2L: Cells were lysed for 2hr at 4°C in NP-40 lysis buffer (1% NP-40, 150 mM NaCl, 50 mM Tris-HCl) with 1% protease inhibitor cocktail (Sigma, P8430), 72 hours after transfection. Samples were separated on either 8% or 15% SDS-PAGE gels and transferred to nitrocellulose membranes using a Western blot apparatus (Bio-rad). Running buffer composition was 190 mM Glycine, 25 mM Tris and 4 mM SDS. Transfer buffer composition was 20% methanol, 3.5 mM sodium carbonate, 10 mM sodium bicarbonate. Membranes were blocked with 5% skim milk dissolved with 1% TBST for 1 hour at room temperature. LMAN2 was detected using rabbit anti-VIP36/LMAN2 primary antibody (Sigma-Aldrich: SAB4200233). LMAN1 was detected using mouse monoclonal anti-LMAN1 primary (ThermoFisher OTI1A8, cat#: MA5-25345). LMAN2L was detected using rabbit polyclonal anti-LMAN2L primary antibody (ThermoFisher cat# PA5-55355). β-actin was detected as the loading control. Chemiluminescence was detected using SuperSignal West Femto Max Sensitivity Substrate (Thermo Fisher Scientific) and a FluorChem SP gel imager (Alpha Innotech).

## RESULTS

### Extrinsic redox sensitive modulation of Kv1.2

Several groups have described highly variable gating of Kv1.2, along with powerful modulation by extracellular redox potential (Grissmer *et al*., 1994; Rezazadeh *et al*., 2007; Baronas *et al*., 2015, 2017). Under ambient redox conditions, Kv1.2 channels expressed in cell lines exhibit wide variation of the voltage-dependence of activation. For example, we observe considerable cell-to-cell variability of V_1/2_ (between ∼−10 mV and 50 mV) when Kv1.2 is expressed in several commonly used cell lines (CHO, L(tk-)) in ambient redox. However, in mild reducing conditions generated by extracellular application of DTT or the membrane impermeant reducing agent TCEP, Kv1.2 channels gate more uniformly and exhibit a prominent depolarizing shift of the activation V_1/2_ to ∼60 mV (Fig. 1A)(Baronas *et al*., 2017). The slow activation properties arising from reducing conditions also lead to a use-dependent activation (UDA) phenotype, in which repetitive trains of depolarizations relieve the inhibitory effect of DTT or TCEP and lead to prominent progressive current enhancement. This property is highly variable under ambient redox conditions, but becomes prominent and consistent in mild reducing conditions (eg. DTT concentrations of 10 μM or higher, Fig. 1B). Due to the extreme variability of these features in different expression systems, and our previous demonstration that these effects are conserved after mutation of endogenous cysteines in Kv1.2 (Baronas *et al*., 2017), we proposed that Kv1.2 gating is influenced by an extrinsic redox-sensitive factor such as a regulatory protein or lipid. A previous investigation of candidate regulators identified by mass spectrometry of Kv1.2 immunoprecipitates led to the identification of Slc7a5 as a powerful regulator of gating and expression in Kv1.1 and Kv1.2 (Baronas *et al*., 2018; Lamothe *et al*., 2020). However, gating effects of Slc7a5 are distinct from the redox-mediated changes in activation kinetics illustrated in Figure 1, suggesting the involvement of other unknown regulatory proteins.

**Figure 1.**
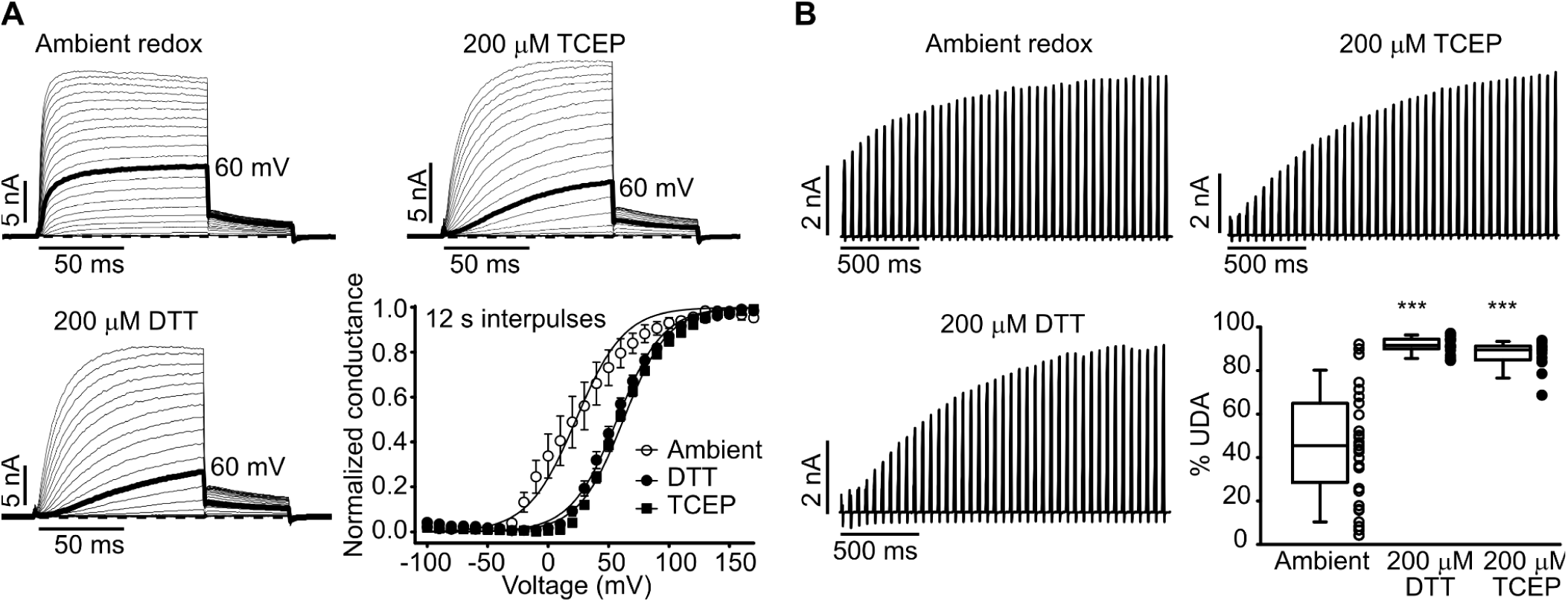
Redox-dependent modulation of Kv1.2. (A) Exemplar current traces and conductance voltage-relationships for Kv1.2 channels. Channels were transiently expressed in LM fibroblasts and pulsed for 100 ms between -100 and 190 mV with 12 s interpulses in ambient redox or indicated redox conditions (200 μM DTT or 200 μM TCEP). Tail currents at -30 mV were analyzed to determine normalized conductance at different prepulse voltages (lower right), and fit with a Boltzmann function (Ambient redox V_1/2_ = 23.1 ± 7.3 mV [mean ± SEM], *k* = 21.2 ± 1.5, n = 9; 200 μM DTT V_1/2_ = 56.9 ± 2.7 mV, *k* = 19.3 ± 0.6, n = 5, 200 μM TCEP V_1/2_ = 62.1 ± 1.9 mV, *k* = 18.8 ± 0.4, n = 8). (B) Exemplar traces and use-dependent activation (% UDA), measured by pulsing repetitively for 10 ms (20 Hz) to 60 mV, from a holding voltage of -80 mV (Ambient redox, n = 39; DTT, n = 17; TCEP, n = 17). A nonparametric Kruskal-Wallis ANOVA on ranks test, followed by a multiple comparison (Dunn’s) post hoc test was used to compare redox conditions (DTT and TCEP) against ambient redox (***p<0.001). Under ambient conditions, use-dependent activation varies considerably (lower right panel), but is markedly enhanced in the presence of extracellularly applied reducing agents.

### Redox-sensitive Kv1.2 modulation in DRG neurons

We previously reported use-dependent activation of Kv1.2-containing channels in dissociated hippocampal neurons, isolated by subtraction of tityustoxin (TsTX)-sensitive currents. However, the redox sensitivity of Kv1.2 gating was not recognized at the time that research was performed (Baronas *et al*., 2015). Prominent currents attributed to Kv1.2-containing channels are also apparent in primary DRG neurons (Zhang *et al*., 2021), where they may influence excitability and pain after nerve-injury (Everill and Kocsis, 1999; Kim *et al*., 2002; Zhao *et al*., 2013, 2017; Fan *et al*., 2014; Laumet *et al*., 2015; Liang *et al*., 2016). We similarly observe Kv1.2-mediated currents in primary DRG cultures, based on a TsTX-subtraction approach (Fig. 2A), and these native currents exhibit prominent redox-sensitive gating (Fig. 2B-E). In patch clamp recordings in ambient redox, TsTx-sensitive currents exhibit variable activation kinetics, quantified here as the rise times between 20% and 80% of peak current (Fig. 2D). However, TsTX-sensitive currents in 200 μM TCEP exhibit prominent slow activation and a rightward shifted V_1/2_ of activation (Fig. 2D-E). As an important control and perhaps of general interest, TsTX retains its activity against Kv1.2 in reducing conditions (Supplementary Fig. 1) - we assessed this because we had concerns that TCEP or other reducing agents could disrupt essential disulfide bonds in TsTX. Lastly, redox-sensitive slow gating of Kv1.2 likely influences excitability of spontaneously firing DRG neurons (Fig. 2F,G), since application of TCEP markedly enhances the spontaneous firing rate, to a comparable extent as channel inhibition by TsTX.

**Figure 2.**
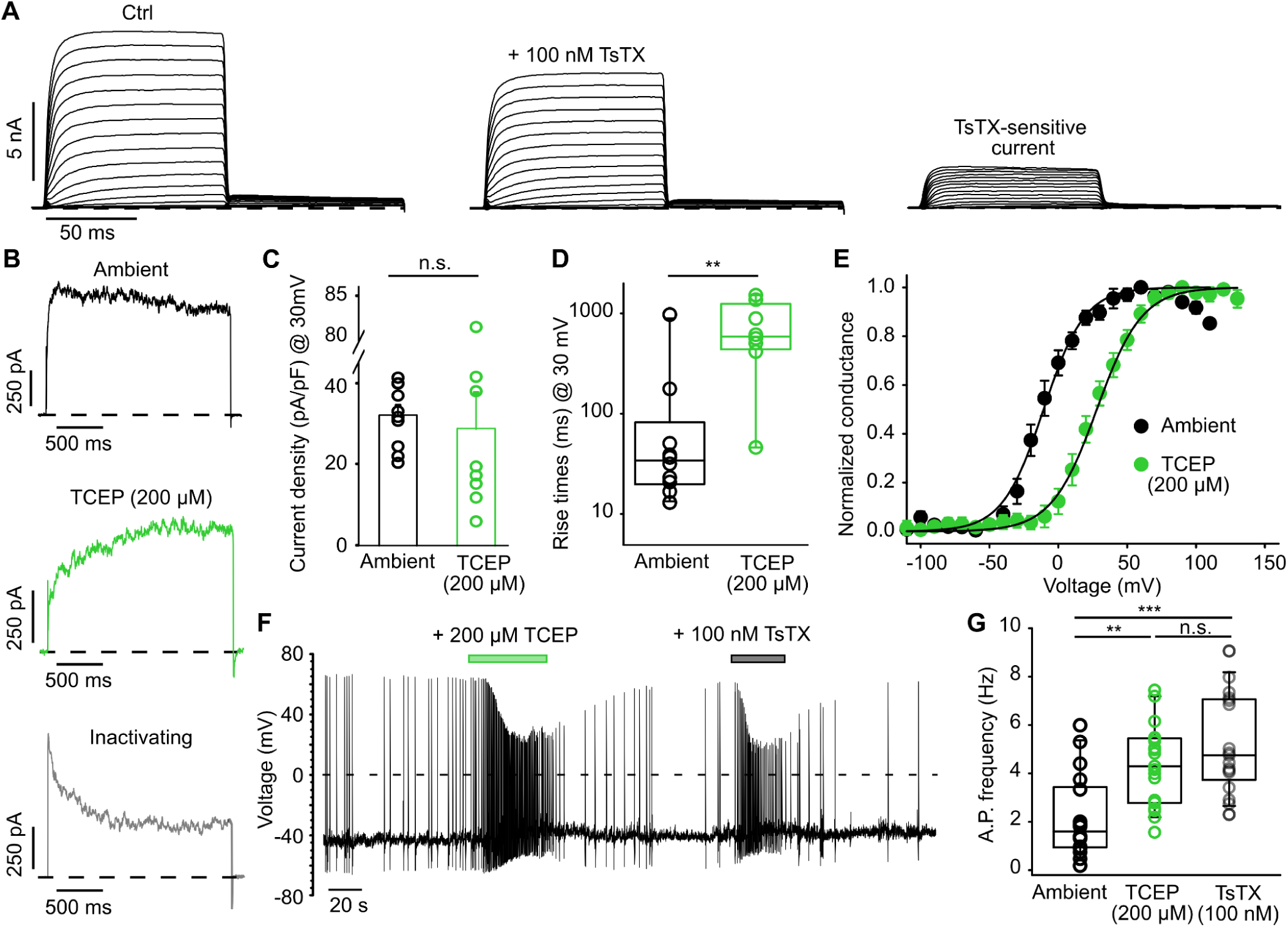
Redox sensitive Kv1.2-containing currents in DRG neurons. (A) Toxin subtraction with 100 nM tityustoxin (TsTX) was used to determine the contribution and properties of Kv1.2-containing currents in DRG neurons of varying size, 1-3 days post-isolation. Cells were held at -100 mV then pulsed between -130 mV and 130 mV (100 ms, 10 mV steps) before stepping to -30 mV (100 ms) in the absence then presence of 100 nM TsTX to assess the toxin-sensitive current by subtraction. An average of 25.5 ± 3.7% [mean ± SEM] of TsTX-subtracted current was observed from total isolated Kv currents. (B) Representative activation traces of TsTX-sensitive currents during a 2 s depolarizing pulse to 30 mV, in ambient redox conditions or in 200 μM TCEP. Occasionally, TsTX-sensitive currents exhibited some degree of inactivation (∼30% of cells), but inactivating currents were rarely observed in TCEP conditions. (C,D) Properties of TsTX-sensitive currents (current density and rise times from 20-80% of peak at 30 mV of a 2 s depolarizing pulse) were determined in ambient redox (n = 10) and 200 μM TCEP conditions (n = 8). A two-tailed Student’s t-test was used to compare TsTX-sensitive current density or activation rise times at 30 mV in ambient redox versus 200 µM TCEP (**p<0.01). (E) Tail currents at -30 mV of TsTX-sensitive currents were analyzed to determine normalized conductance at different prepulse voltages (-130 mV to 130 mV, 10 mV increments) and fit with a Boltzmann function (ambient redox V_1/2_ = -10.7 ± 4.5 mV [mean ± SEM], *k* = 17.3 ± 2.0, n = 11; 200 μM TCEP V_1/2_ = 27.9 ± 4.6 mV, *k* = 17.7 ± 1.5, n = 16. (F) Exemplar records of spontaneous action potential firing of a DRG neuron (0 pA holding current), under ambient redox conditions, and application of 200 μM TCEP or + 100 nM TsTX. (F) Frequency (Hz) of action potential firing was assessed at 0 pA holding current in ambient redox, + 200 μM TCEP and + 100 nM TsTX conditions (n = 17-19). A nonparametric Kruskal-Wallis ANOVA on ranks test, followed by a multiple comparison (Dunn’s) post hoc test was used to compare between 200 μM TCEP, 100 nM TsTX and ambient redox conditions (***p<0.001, **p<0.01).

### LMAN2 is a candidate regulator of use-dependent Kv1.2 gating

To explore mechanisms underlying variability of Kv1.2 gating, we investigated candidate regulatory proteins identified by mass spectrometry profiling of Kv1.2 immunoprecipitates (Baronas *et al*., 2018). We quantified gating properties of Kv1.2 in individual cells using the ‘use-dependent activation’ parameter, which describes the % activation between the first pulse and peak pulse in a train of depolarizations (60 mV for 10 ms, 20 Hz frequency). As described previously, Kv1.2 exhibits highly variable use-dependence. In reducing conditions, ‘use-dependent’ gating properties are prominent and consistent, whereas in ambient extracellular redox conditions, use-dependent gating is far more variable. We co-expressed and characterized 52 candidate regulators with Kv1.2 and assessed use-dependent activation in ambient redox (Fig. 3A). Most candidate regulatory proteins had high variability with no consistent effect on Kv1.2 activation gating. However, co-expression with LMAN2 caused a consistent bias towards strong use-dependent activation (Fig. 3B,C). When co-expressed with LMAN2, Kv1.2 exhibited mean use-dependent activation of 86%, and no cell exhibited less than 62% use-dependent activation.

**Figure 3.**
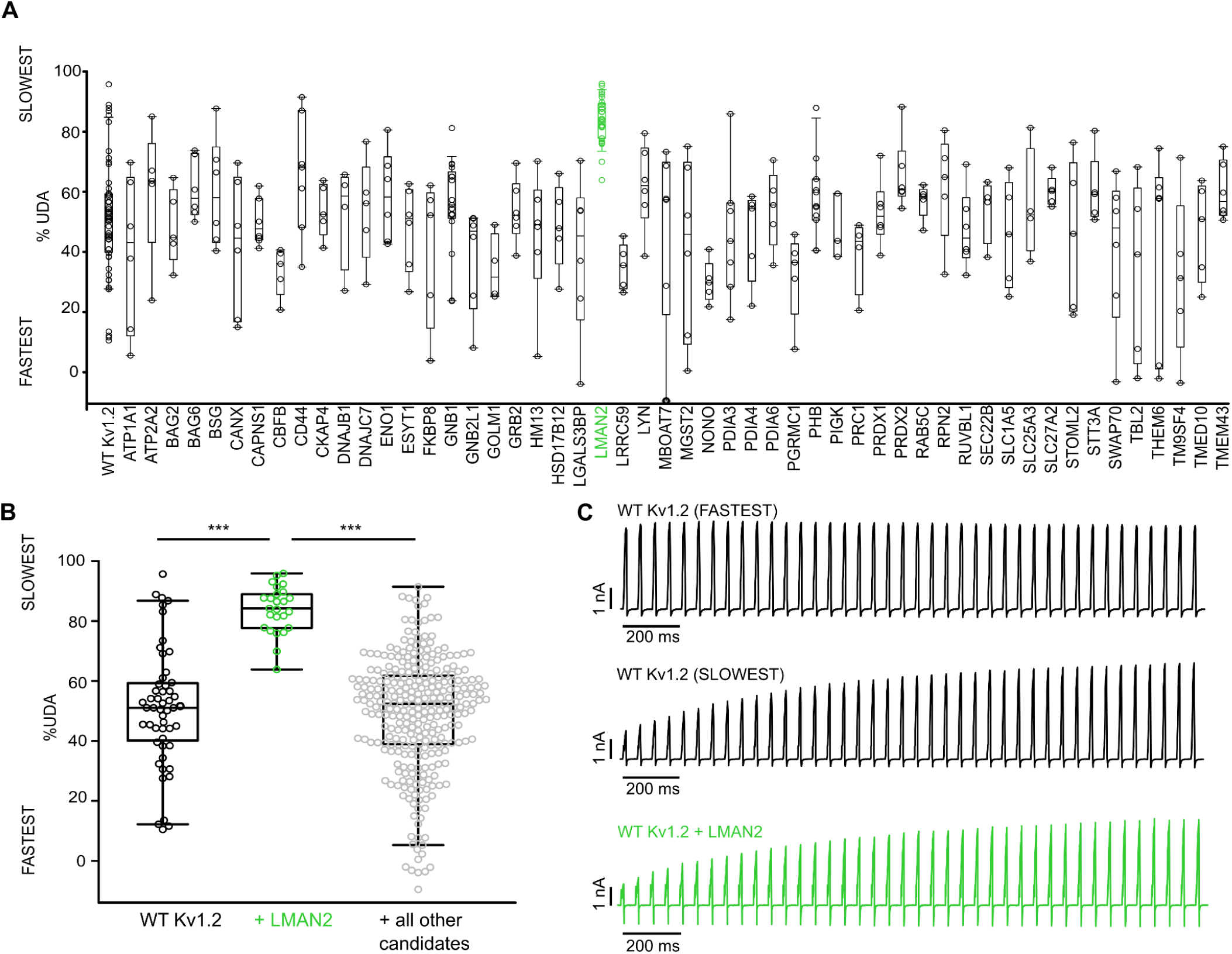
Screening of candidate regulators of Kv1.2 use-dependent activation. **(A)** use-dependent activation (% UDA) was measured by patch clamp recordings of 5 or more LM fibroblasts transfected with Kv1.2 and each of the indicated candidate genes (identified by mass spectrometry analysis of Kv1.2 Immunoprecipitated proteins and organized in alphabetical/numerical order). Co-expression with LMAN2 biases use-dependence of Kv1.2 towards a slow gating phenotype. (B) Distribution of use-dependence observed from Kv1.2-co-transfected with LMAN2 (green), relative to Kv1.2 alone (black), or with all other candidate regulators (gray). A nonparametric Kruskal-Wallis ANOVA on ranks test, followed by a multiple comparison (Dunn’s) post hoc test was used to compare Kv1.2 + LMAN2 against WT Kv1.2 and Kv1.2 + all other candidates (***p<0.001). (C) Exemplar traces illustrate the extreme ends (FASTEST & SLOWEST) of variable Kv1.2 use-dependent activation, versus predominantly strong use-dependence when co-expressed with LMAN2.

### Characterization of Kv1.2 modulation by LMAN2

We characterized the effects of LMAN2 modulation of Kv1.2 in more detail. We confirmed that LMAN2 co-expression biases cells towards a prominent slow use-dependent behavior (Fig. 4A-C). We also quantified LMAN2 effects on activation kinetics using the 20-80% rise time, illustrating marked deceleration of Kv1.2 activation when co-expressed with LMAN2 (Fig. 4B,D,E). Lastly, co-expression with LMAN2 causes a prominent depolarizing shift of activation (V_1/2_), comparable to the saturating effect of reducing agents such as DTT or TCEP (compare with Fig. 1). It is noteworthy that previous work has demonstrated that slow activation of Kv1.2 can be transiently relieved by strong repetitive depolarizations, which shifts channels into a potentiated/facilitated mode (Rezazadeh *et al*., 2007; Baronas *et al*., 2017). This pulse-dependent potentiation of gating is also apparent for LMAN2-modulated currents. That is, channels co-expressed with LMAN2 exhibit slow activation kinetics with a depolarized V_1/2_ (Fig. 4 F,G). However, by adding a conditioning prepulse (60 mV) before each voltage step, LMAN2 inhibition is transiently relieved and channels are shifted into a ‘permissive’ potentiated mode with faster kinetics, and a more hyperpolarized V_1/2_ (Fig. 4 F,G). To observe maximal LMAN2 or TCEP/DTT modulation of Kv1.2 gating, relatively long (∼10-12 s) interpulse intervals are used to allow channels to recover their slow gating mode. In contrast, for generation of potentiated conductance-voltage relations we include a prepulse, 100 ms before each voltage step. These results reinforce that Kv1.2 use-dependent activation reflects a dynamically reversible shift between slow and potentiated gating modes. This balance was previously shown to be strongly biased towards slow gating by reducing agents, and our findings now demonstrate that LMAN2 co-expression mimics the effects of reducing conditions.

**Figure 4.**
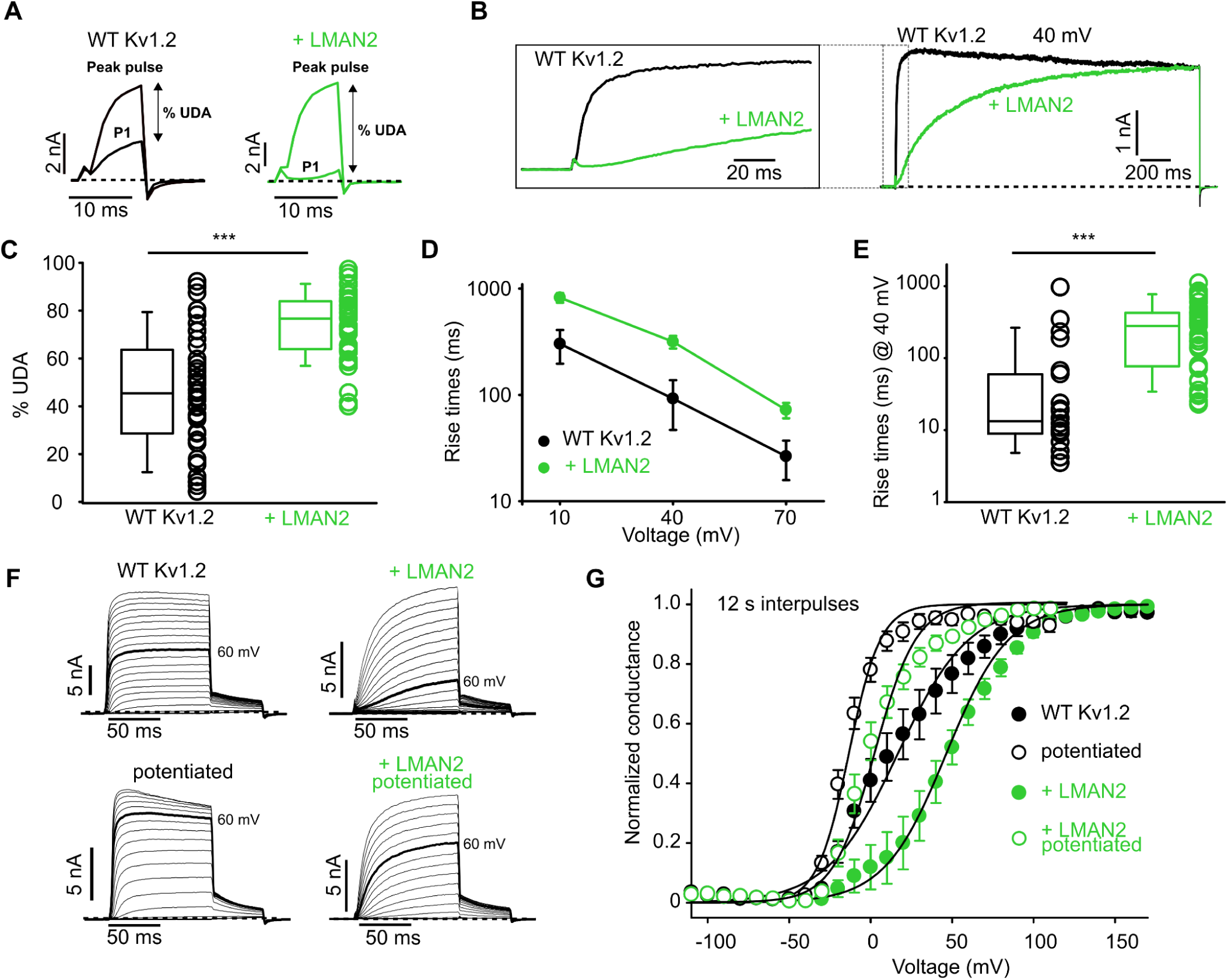
Biophysical features of Kv1.2 co-expressed with LMAN2. (A,C) Measurement of use-dependent activation (UDA). Use-dependence was assessed by pulsing from a holding voltage of -80 mV to a pulse voltage of 60 mV for 10 ms (20 Hz). (C) % UDA was assessed in multiple cells (n = 39 for WT Kv1.2, and n = 37 for Kv1.2 + LMAN2). A two-tailed Welch’s t-test was used to compare the % UDA of WT Kv1.2 versus WT Kv1.2 + LMAN2 (***p<0.001). (B) Kinetics of activation of Kv1.2 co-expressed with LMAN2. Exemplar currents depict typical kinetics of activation of Kv1.2 expressed alone or with LMAN2, elicited by a voltage-step to 40 mV. The inset expanded panel highlights a prominent delay of Kv1.2 activation when co-expressed with LMAN2. (D,E) Kinetics of activation were quantified by measuring the rise times from 20-80% of peak current (n = 22 for WT Kv1.2, n = 41 for Kv1.2 + LMAN2). A Mann-Whitney U-test was used to compare the rise times at 40 mV between WT Kv1.2 and WT Kv1.2 + LMAN2 (***p<0.001). (F,G) Conductance-voltage relationships were determined for Kv1.2 alone, or co-expressed with LMAN2, stepping between -100 and 190 mV in 10 mV steps (100 ms). ‘Potentiated’ refers to inclusion of a 100 ms, 60 mV prepulse shortly before (100 ms) each sweep of the conductance-voltage protocol (WT Kv1.2 V_1/2_ = 16.7 ± 4.6 mV [mean ± SEM], *k* = 22.6 ± 1.0, n = 14; ‘potentiated’ V_1/2_ = -13.4 ± 2.5 mV, *k* = 10.8 ± 1.4, n = 8 ; WT Kv1.2 + LMAN2 V_1/2_ = 46.4 ± 6.8 *k* = 19.1 ± 2.3, n = 7; potentiated LMAN2 V_1/2_ = 2.4 ± 3.7, *k* = 14.2 ± 1.3, n = 11).

### Knockdown and rescue of LMAN2

Since Kv1.2 use-dependent gating is highly variable in ambient redox conditions, we suspect that endogenous levels of one or more mediators (potentially including LMAN2) influence Kv1.2. We tested the influence of endogenous LMAN2 using shRNA knockdown in LM cells, using lentiviral delivery and puromycin selection to maintain stable shRNA expression. We isolated individual clonal cells by serial dilutions, and selected a cell line with prominent knockdown of LMAN2, confirmed with Western blot (Fig. 5A,B). Kv1.2 expressed in LMAN2 shRNA cells exhibited significantly weakened use-dependent activation properties and less variability, compared to cells expressing a scramble shRNA (Fig. 5C,D). Furthermore, knockdown of LMAN2 weakened responsiveness to reducing conditions. For example, application of 30 μM DTT elicits prominent use-dependent activation in scramble shRNA LM cells, but only weak use-dependence of Kv1.2 in LMAN2 shRNA LM cells (Fig. 5C,D). Additionally, transfection with human LMAN2 led to increased use-dependent gating in ambient redox, and rescued sensitivity of Kv1.2 to 30 μM DTT (Fig. 5C,D). These findings indicate that LMAN2 is a contributor to redox sensitive use-dependent gating of Kv1.2 observed in mammalian cell lines.

**Figure 5.**
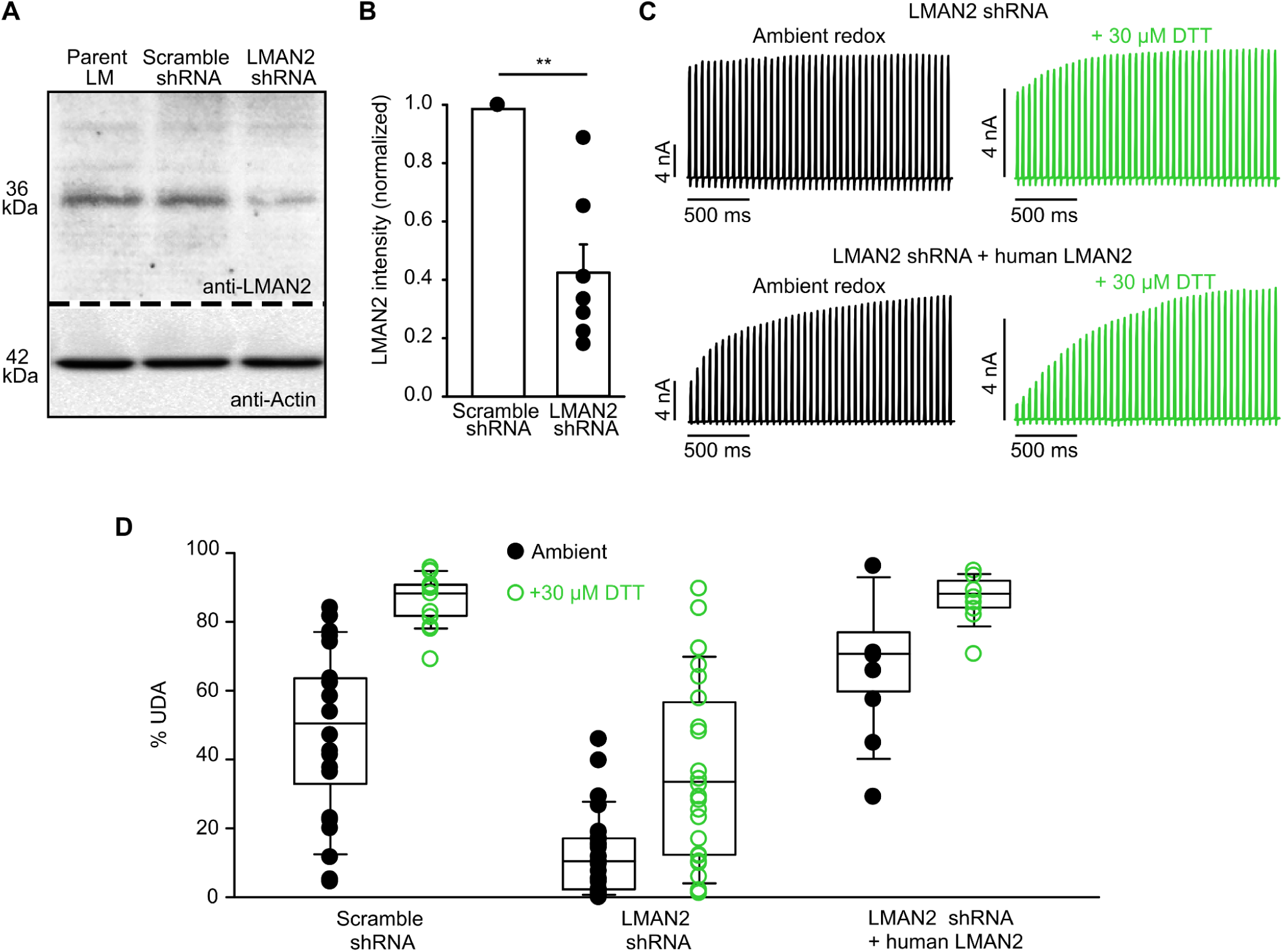
LMAN2 knockdown attenuates redox-sensitive use-dependent activation of Kv1.2. (A, B) An LMAN2 knockdown LM fibroblast cell line was established by lentiviral transduction and stable expression of an LMAN2-targeted shRNA, resulting in an average of ∼60% knockdown based on anti-LMAN2 western blots (n = 7) (two tailed Student’s t-test between scrambled shRNA and LMAN2 shRNA, **p<0.01). (C) Exemplar traces depicting use-dependent activation of Kv1.2 in LMAN2-knockdown cells +/- overexpressed human LMAN2, determined in ambient redox and mild reducing conditions (30 μM DTT). (D) Summarized use-dependent activation properties and redox-sensitivity of use-dependent activation (% UDA) of scramble shRNA cells, LMAN2 shRNA cells, and rescue of redox modulation by human LMAN2. For Scramble shRNA: (Ambient, n = 26) (+30 μM DTT, n = 19); for LMAN2 shRNA: (Ambient, n = 31) (+30 μM DTT, n = 30); for LMAN2 shRNA + human LMAN2: (Ambient, n = 12) (+30 μM DTT, n = 12).

### LMAN2 overexpression leads to enhanced redox sensitivity

LMAN2 co-expression mimics the functional outcomes of extracellular reducing conditions, raising the possibility that LMAN2 is involved in the cellular mechanism underlying redox sensitivity of Kv1.2. Thus, we further tested if LMAN2 could influence Kv1.2 responses to reducing conditions. In reducing conditions (200 μM DTT in Fig. 6), co-expression with LMAN2 does not further enhance use-dependent activation and has only modest effects on activation kinetics or V_1/2_ (Fig. 6A-C). Thus, co-expression with LMAN2 does not have an additive effect above the gating effects arising from DTT alone. These findings suggest that LMAN2 influences the endogenous mechanism of redox sensitivity, rather than operating by a distinct mechanism. To test this further, we determined the effects of LMAN2 on the concentration response to DTT. For these experiments we used Chinese Hamster Ovary (CHO) cells because we have observed they exhibit moderate use-dependent activation under ambient redox conditions, so effects of LMAN2 or reducing agents can be more apparent than in LM cells. We determined an EC_50_ of 1.8 μM DTT, for enhancement of use-dependent activation in Kv1.2 expressed alone, whereas LMAN2 sensitized channels to DTT roughly 10-fold, leading to an EC_50_ of 0.16 μM (Fig. 6D). This sensitization to DTT is also apparent in terms of dynamics of use-dependent activation (Fig. 6E). In this experiment, we measured potentiation of current during the standard train of depolarizations (20 Hz) to elicit use-dependent activation. When co-expressed with LMAN2, channels are sensitized to low concentrations (200 nM) of DTT, thereby exhibiting slower potentiation, and requiring more pulses to reach maximal potentiation. However, at saturating concentrations of DTT (200 μM), LMAN2 no longer causes this sensitization (Fig. 6E, inset), consistent with DTT and LMAN2 acting via overlapping mechanisms.

**Figure 6.**
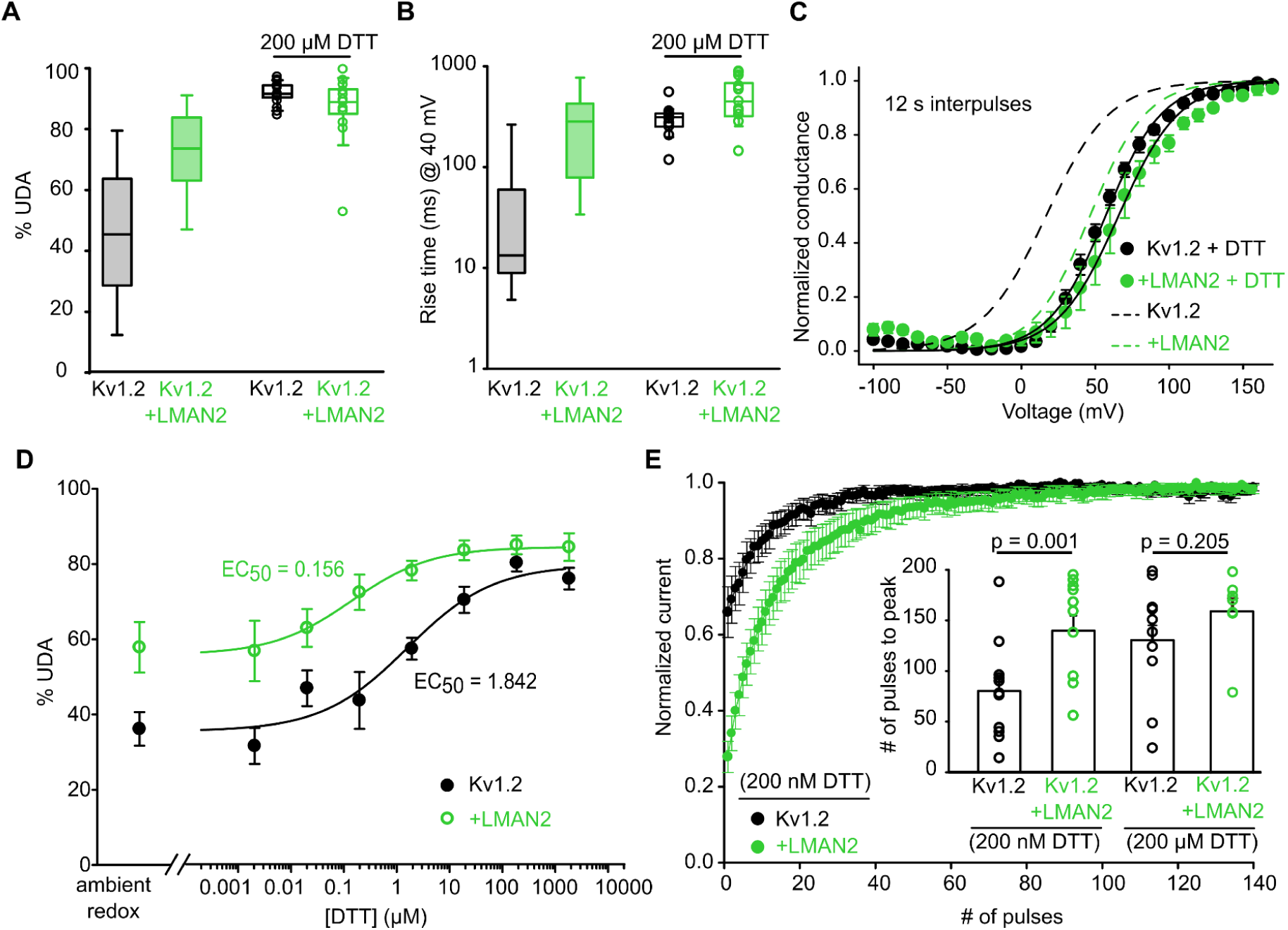
LMAN2 overexpression sensitizes CHO cells to reducing agents and use-dependent activation. (A,B) Use-dependent activation (% UDA, n = 13-17) and rise times (20-80% at 40 mV, n = 13-14) of WT Kv1.2 +/- LMAN2 were measured in LM cells in the indicated conditions (Boxes shaded in grey and green are superimposed from Figure 4C and E for comparison). DTT treatment, or co-expression with LMAN2 enhances Kv1.2 use-dependent activation and decelerates rise times but the effect is not additive. (C) Conductance-voltage relationships were measured as described in Figure 1A. (Kv1.2 + 200 μM DTT V_1/2_ = 56.9 ± 2.7 mV [mean ± SEM], *k* = 19.3 ± 0.6, n = 5; Kv1.2 + LMAN2 + 200 μM DTT V_1/2_ = 65.7 ± 7.1 mV, *k* = 20.8 ± 0.8 mV, n = 7). (Dotted lines are duplicated from Figure 4G). (D) Use-dependent activation (% UDA) was assessed over a range of DTT concentrations for WT Kv1.2 alone or co-expressed with LMAN2 in CHO cells (for Kv1.2 alone, n = 7-25; for Kv1.2 + LMAN2, n = 4-13). LMAN2 co-expression leads to more prominent use-dependent activation in ambient redox, and a sensitization to lower concentrations of DTT. (E) Pulse-by-pulse development of current potentiation during repetitive trains of depolarizations (20 Hz), in low (200 nM) and high (200 μM) concentrations of DTT was measured and summarized based on the number of depolarizations required to reach the peak current. For 200 nM DTT: (Kv1.2, n = 11) (Kv1.2 + LMAN2, n = 12); for (200 μM DTT: (Kv1.2, n = 10) (Kv1.2 + LMAN2, n = 7). A two-tailed Student’s t-test was used to compare Kv1.2 + LMAN2 versus Kv1.2 alone in 200 nM or 200 µM DTT; p-values are denoted above the respective bar graphs.

### Mapping structural determinants of LMAN2 and redox sensitivity

Previous reports investigated channel segments that control susceptibility to slow gating, and identified residue T252 in the Kv1.2 S2-S3 linker as a critical position. Kv1.2 [T252R] channels (and certain other mutations at this position) weaken use-dependence and the slow gating mode of Kv1.2 (Rezazadeh *et al*., 2007; Baronas *et al*., 2017). We have now recognized additional complexity to this effect, as the T252R mutation does not completely abolish use-dependent gating properties. Instead, some use-dependence and gating shift clearly persists for T252R channels in reducing conditions. (Fig. 7A-C). Similarly, the T252R mutant channel exhibits weakened, but not abolished, sensitivity to LMAN2 (Fig. 7A-C), with markedly reduced use-dependence, faster rise times of activation, and an incomplete bimodal V_1/2_ shift relative to WT Kv1.2. However, chimeric substitution of the S2-S3 linker of Kv1.2 with Kv1.5, comprising just one additional mutation (T252R + F251S) is sufficient to completely abolish use-dependent gating phenotype and modulation by DTT and LMAN2 (Fig. 7D-G-J). The effect of the Kv1.2[F251S] mutation alone is not dramatic, as these channels exhibit use-dependence and retain sensitivity to both LMAN2 and DTT (Fig. D-G). Other chimeras tested, which switched either the S1, S1-S2 linker, or S2 segments of Kv1.2 with sequence from Kv1.5, retained sensitivity to both LMAN2 and DTT (Supplementary Fig. 2A-C). Swapping the entire S1-S3 segment with Kv1.5 sequence completely abolished the sensitivity to both LMAN2 and DTT (Supplementary Fig. 2D). Since redox modulation and LMAN2 modulation have overlapping structural determinants localized to the S2-S3 linker, we suspect these treatments act by a shared mechanism.

**Figure 7.**
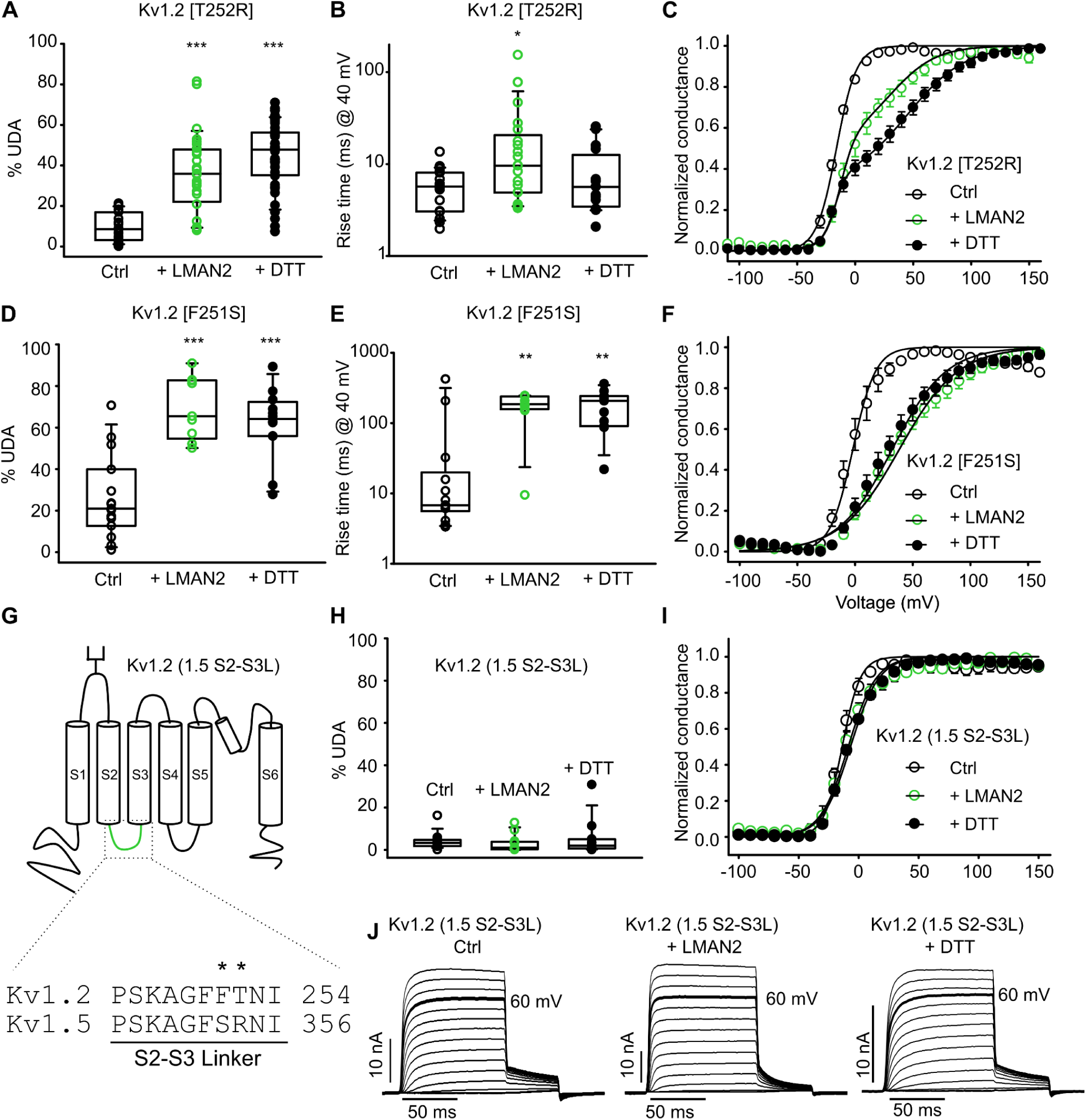
Overlapping determinants of LMAN2 and reducing agent sensitivity of Kv1.2. (A-C) % UDA, rise times of activation (20-80% at 40 mV), and conductance-voltage relationships were determined for Kv1.2 [T252R] (% UDA: Ctrl n = 26, LMAN2 n = 27, 200 μM DTT n = 29) (Rise times: Ctrl n = 21, + LMAN2 n = 20, + 200 μM DTT n = 15) in ambient redox, 200 μM DTT, or co-expressed with LMAN2, as indicated. A nonparametric Kruskal-Wallis ANOVA on ranks test, followed by a multiple comparison (Dunn’s) post hoc test was used to compare Kv1.2[T252R] in ambient redox with 200 μM DTT or LMAN2 conditions in A & B (***p<0.001) (*p<0.05). Kv1.2[T252R]: V_1/2_ = -16.2 ± 0.9 mV [mean ± SEM]; *k* = 8.6 ± 0.4 mV; n = 10; Kv1.2[T252R] + LMAN2: V_1/2_a = -13.0 mV; *k* = 6.6 mV; V_1/2_b = 31.4 mV; *k* = 19.4 mV; n = 9; Kv1.2[T252R] + 200 μM DTT: V_1/2_a = -17.8 mV; *k* = 5.3 mV; V_1/2_b = 45.1 mV; *k* = 24.8 mV; n = 16 (D-F) % UDA, rise times of activation (20-80% at 40 mV), and conductance-voltage relationships were determined for Kv1.2[F251S] (%UDA: Ctrl n = 15, + LMAN2 n = 9, + 200 μM DTT n = 12) (Rise times: Ctrl n = 14, + LMAN2 n = 10, + 200 μM DTT n = 11) in ambient redox, 200 μM DTT, or co-expressed with LMAN2, as indicated. A One-way ANOVA followed by a Dunnett’s post hoc test (D) and a nonparametric Kruskal-Wallis ANOVA on ranks test, followed by a multiple comparison (Dunn’s) post hoc test (E) was used to compare Kv1.2[F251S] in ambient redox with 200 μM DTT or LMAN2 conditions (***p<0.001)(**p<0.01). Kv1.2[F251S]: V_1/2_ = -1.9 ± 3.3 mV [mean ± SEM]; *k* = 10.3 ± 0.8 mV n = 10; Kv1.2[F251S] + LMAN2: V_1/2_ = 38.3 ± 5.1 mV; *k* = 26.3 ± 2.2 mV n = 6; + 200 μM DTT: _V1/2_ = 32.2 ± 4.6 mV; *k* = 24.2 ± 1.9 mV n = 7. (G) Cartoon illustrating the transmembrane topology and location of positions F251 and T252 in the intracellular S2-S3 linker of Kv1.2. (H,I,J) % UDA, conductance-voltage relationships and representative traces for Kv1.2 (1.5 S2-3L) channels in control, 200 μM DTT, or LMAN2 conditions. (%UDA: Ctrl n = 11, + LMAN2 n = 11, + 200 μM DTT n = 10); Kv1.2(1.5 S2-S3L) V_1/2_ = -13.7 ± 1.6 mV; *k* = 7.9 ± 0.8 mV n = 5; + LMAN2: V_1/2_ = -8.7 ± 3.3 mV; *k* =12.6 ± 1.5 mV n = 7; + 200 μM DTT: V_1/2_ = -6.8 ± 1.1 mV; *k* = 11.9 ± 0.3 mV n = 8.

### Cell surface localization of LMAN2 is required for Kv1.2 modulation

LMAN2 is a single-pass transmembrane protein with a large extracellular N-terminal lectin domain and a short intracellular C-terminus. There are varied reports on localization of LMAN2, which may depend on expression level and cell type, but several studies have demonstrated cell surface localization of some fraction of LMAN2 (Shirakabe *et al*., 2011; Hua *et al*., 2024). We used cell surface biotinylation to test whether LMAN2 and other members of the L-type transmembrane lectin family (LMAN1 and LMAN2L) can reach the plasma membrane (Fig. 8A-C) and influence Kv1.2 gating (Fig. 8E&F). Importantly, the C-termini of LMAN2 and other related transmembrane lectins have sequence differences that may influence cellular localization (Fig. 8D). In particular, LMAN1 (C-terminal KKFF) and LMAN2L (C-terminal RKR) have strong ER retention/retrieval signals that are absent in LMAN2 (Fig. 8D) (Arar *et al*., 1995; Itin *et al*., 1995; Hauri *et al*., 2000; Shirakabe *et al*., 2011; Qin *et al*., 2012). Consistent with this and prior work, neither LMAN1 or LMAN2L exhibited detectable cell surface localization (Fig. 8A,B)(Neve *et al*., 2003; Nufer *et al*., 2003). In contrast, a significant fraction of EGFP-LMAN2 (N-terminal tag) reached the cell surface (Fig. 8C), and this was abolished by introducing the ‘KKAA’ ER retention signal to the C-terminus of LMAN2 (Fig. 8C)(Shirakabe *et al*., 2011). It is noteworthy that Western blots for EGFP-LMAN2 (N-terminal tag) consistently exhibited two bands, which is likely due to cellular proteolytic processing of the N-terminal tag during biogenesis or sample preparation (Fig. 8C). These patterns of lectin localization were consistent with electrophysiological effects in Kv1.2 (Fig. 8E,F). Kv1.2 co-expression with untagged LMAN2 (Fig. 8E,F) or N-terminally tagged EGFP-LMAN2 (Figs.3,4) consistently led to slow activation properties and use-dependence. However, this was not observed for intracellularly localized LMAN1, LMAN2L or LMAN2 (KKAA) (Fig. 8E-F). Additionally, a C-terminal EGFP-tagged version of LMAN2 (LMAN2-EGFP) was also insufficient to promote slow gating of Kv1.2 (Fig. 8E-F), and was not detectable in the surface biotinylated protein fraction (data not shown). This was consistent with early work using C-terminally tagged LMAN2 constructs that reported intracellular retention of LMAN2 (Fiedler *et al*., 1994), highlighting that the addition (and location) of an epitope or fluorescent protein tag can prominently influence LMAN2 localization. Taken together, these findings indicate that cell surface localization of LMAN2 is essential for modulation of Kv1.2 gating.

**Figure 8.**
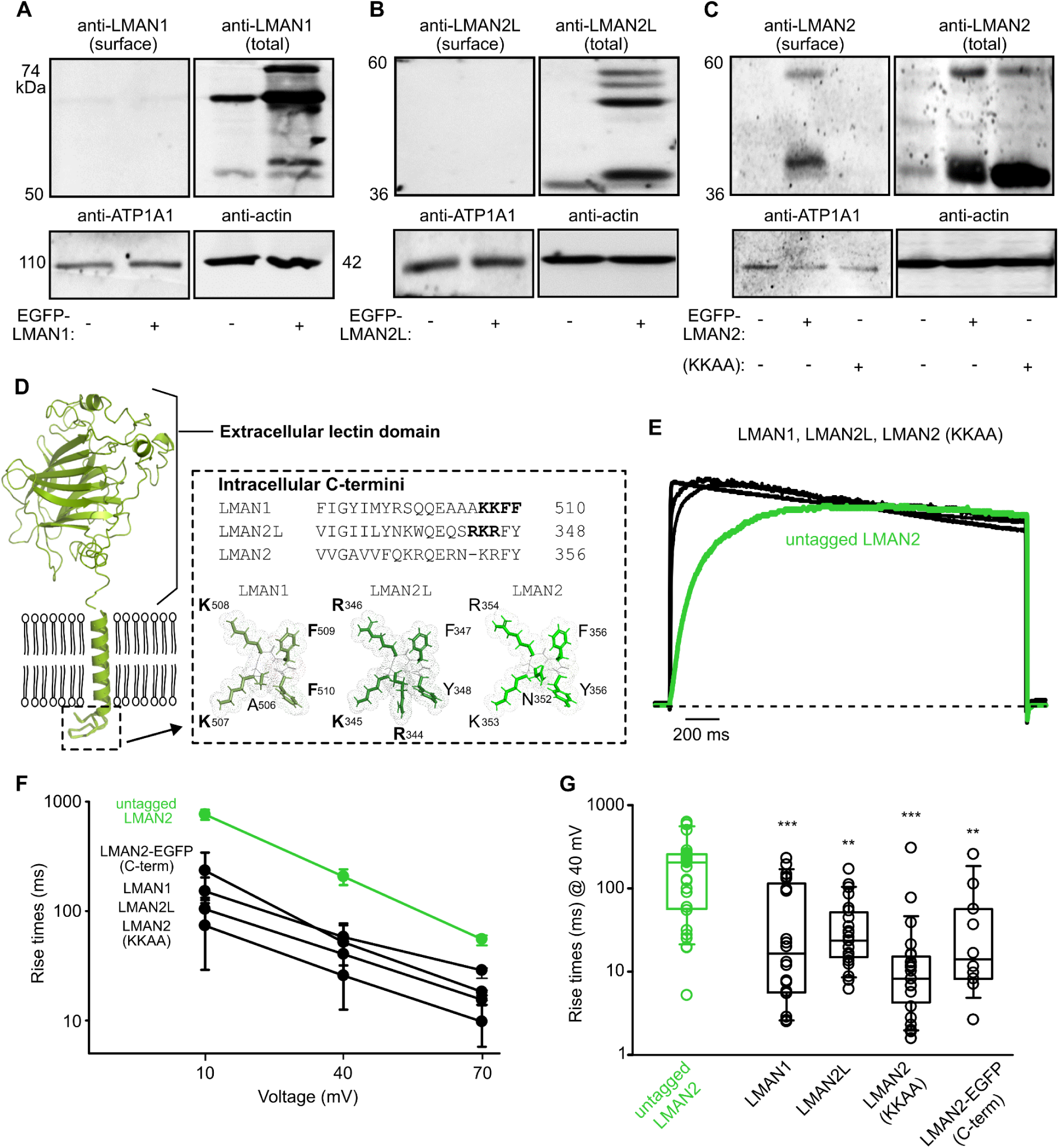
Kv1.2 modulation requires cell surface expression of LMAN2. (A-C) Cell surface biotinylated fractions and total cell lysates isolated from HEK293 cells transfected with various LMAN proteins. EGFP-LMAN1(A), EGFP-LMAN2L(B), and EGFP-LMAN2 [KKAA](C) fail to generate prominent localization at the cell surface, whereas EGFP-LMAN2(C) is detectable at the cell surface. (D) Transmembrane topology of LMAN2 and relatives, highlighting a large globular extracellular lectin domain, a single-pass transmembrane domain, and variable C-termini. Sequences of the final 19 amino acids of LMAN1, LMAN2, and LMAN2L are shown, with ER retention/retrieval signals in LMAN1 and LMAN2L indicated by bold lettering. AlphaFold Predicted protein structures (LMAN1: AF-P49257-F1; LMAN2L: AF-Q9H0V9-F1; LMAN2: AF-Q12907-F1) depict the ER retention/retrieval signal amino acids in LMAN1 and LMAN2L (bold lettering) but not LMAN2. (E) Exemplar current traces elicited at 40 mV for 2 s for Kv1.2 co-expressed with untagged LMAN2, LMAN1, LMAN2L, LMAN2 [KKAA] (introduced ER retention signal) or LMAN2-EGFP (C-term), as indicated. (F,G) Rise times (20%-80% of peak current) were assessed for Kv1.2 co-expressed with each LMAN subtype (n = 28 for untagged LMAN2, n = 20 for LMAN1, n = 24 for LMAN2L, n = 24 for LMAN2 (KKAA), n = 10 for LMAN2-EGFP (C-term). A nonparametric Kruskal-Wallis ANOVA on ranks test, followed by a multiple comparison (Dunn’s) post hoc test was used to compare Kv1.2 + untagged LMAN2 against all other LMAN subtypes (***p<0.001) (**p<0.01). Significant modulation of Kv1.2 is only apparent when co-expressed with LMAN2 (untagged) or EGFP-LMAN2.

## Discussion

Kv1.2 is an essential regulator of neurological function as it influences action potential firing, duration, and morphology. Downregulation or knockout of Kv1.2 leads to generalized seizures and death in rodent models (mice, gerbils), and both gain- or loss-of-function Kv1.2 mutations in humans lead to severe seizures and neurodevelopmental delay (Buckley *et al*., 2001; Brew *et al*., 2007; Lee *et al*., 2009; Syrbe *et al*., 2015; Masnada *et al*., 2017). Another pathological outcome of Kv1.2 dysfunction is neuropathic pain arising from decreased Kv1.2 expression that leads to hyperexcitable primary afferent DRG neurons (Ishikawa *et al*., 1999; Zhao *et al*., 2017; Sun *et al*., 2019), while restoration of Kv1.2 expression in nerve-injured DRG neurons alleviates neuropathic pain (Zhang *et al*., 2021). In this study, we directly measured acute effects of tityustoxin (TsTX)-mediated Kv1.2 inhibition in DRG neurons (Fig. 2), leading to increased action potential firing in current clamp recordings. The TsTX-sensitive Kv currents likely arise from diverse Kv1.2-containing heteromeric channels, as TsTX remains a relatively effective blocker of Kv1.2 co-assembled with other related Kv1 subunits with varying potency. Kv1.1, Kv1.4, or Kv1.5 subunits in combination with Kv1.2 exhibit, ∼80%, ∼25%, and ∼25% inhibition by 100 nM TsTX, respectively (Sheng *et al*., 1993; Wang *et al*., 1993; Werkman *et al*., 1993; Baronas *et al*., 2015). Our findings also indicate that suppression of Kv1.2 (in reducing conditions) makes a significant contribution to redox-dependent changes in DRG firing. Both TsTX and reducing agents cause similar enhancement of firing frequency in isolated DRG neurons, and do not have additive effects (Fig. 2), suggesting that they are converging on the same mechanism of regulation.

Based on our findings in DRG neurons and previous reports in isolated hippocampal neurons, we suspect that redox modulation of Kv1.2 can influence neuronal excitability (Baronas *et al*., 2015)(Fig. 2). The use-dependent activation of Kv1.2 channels has not yet been linked to a specific physiological outcome *in vivo*, but we speculate use-dependent cumulative activation of Kv1.2 could influence action potential firing during bursts of activity. Heterologous and native hippocampal cells exhibit variable Kv1.2-mediated use-dependent activation (Baronas *et al*., 2015). When this modulation is maximized by application of reducing agents (DTT, TCEP), we consistently observe an 80 to 100-fold current increase during a 20 Hz train of depolarizations (Figure 3 and 4), and this dynamic change of Kv1.2-encoded current would likely influence firing, given the powerful effects we have observed after toxin-mediated Kv1.2 inhibition. Also, it is noteworthy that redox-dependent Kv1.2 modulation is prominent in a physiological range of extracellular redox potentials (Baronas *et al*., 2017).

High cell-to-cell variability (and pulse-to-pulse variation) of Kv1.2 activation kinetics were initially reported shortly after Kv1.2 was first cloned (McKinnon, 1989; Grissmer *et al*., 1994). The mechanistic basis for this unique property (Fig. 1) has remained an enigma, although there are clear signs that it is not directly encoded by Kv1.2. The effect varies dramatically between different cell types and even different individual cells expressing the same cDNA clone (Rezazadeh *et al*., 2007; Baronas *et al*., 2015, 2016). Prior work demonstrated a powerful influence of extracellular redox potential, suggesting involvement of an extracellular redox-sensing disulfide bond, however this effect is not mediated by cysteines in Kv1.2 (Baronas *et al*., 2017). Our functional screening approach to test candidate regulatory partners recognized LMAN2 as the strongest protein regulator of Kv1.2 activation kinetics identified thus far (Fig. 3). Diverse evidence supports this finding although there is still uncertainty about the detailed mechanism of LMAN2-mediated channel modulation. For example, the rate and voltage-dependence of Kv1.2 activation are both profoundly shifted when channels are co-expressed with EGFP-LMAN2 or untagged LMAN2, mimicking effects of extracellular reducing agents. Effects of reducing agents and LMAN2 are also not additive (ie. LMAN2 co-expression does not shift channel gating beyond the effects of DTT or TCEP application), suggesting these manipulations are converging on the same regulatory pathway (Fig. 6). The slow gating properties and redox-dependence of Kv1.2 are also blunted by shRNA-mediated knockdown of LMAN2 (Fig. 5).

Our demonstration of the effects of LMAN2 was unexpected because it has not previously been identified as an ion channel regulator. LMAN2 is primarily recognized as a cargo receptor that identifies high-mannose type glycosylation in the secretory pathway and contributes to appropriate cell surface targeting of proteins (Fiedler and Simons, 1994; Fiedler *et al*., 1994; Füllekrug *et al*., 1999; Hara-Kuge *et al*., 2002). Along with other closely related L-type lectins, the structural basis for binding of the lectin domain (topologically lying on the luminal/extracellular side of the membrane) to sugar groups has been determined, along with its role in transporting a few candidate cargo proteins used for model studies, such as alpha-amylase, α1 anti-trypsin and alpha-fetoprotein (Hara-Kuge *et al*., 2004; Kamiya *et al*., 2005, 2008; Reiterer *et al*., 2010; Muranaka *et al*., 2023). There has been some uncertainty about the extent of localization of LMAN2 at the plasma membrane. Some work using antibodies to label endogenous LMAN2 has suggested the absence of this protein at the cell surface, while other work has shown metalloproteinase-dependent shedding of LMAN2 from the cell surface of Raw 264.7 (macrophages), HeLa and HEK293T cells (Füllekrug *et al*., 1999; Hara-Kuge *et al*., 2002; Shirakabe *et al*., 2011; Hua *et al*., 2024). There are suggestions that localization may vary in different cell types and depend on expression level (Yamashita *et al*., 1999; Hauri *et al*., 2000; Schrag *et al*., 2003). In our case, there was a clear requirement for cell surface localization of LMAN2 in terms of Kv1.2 modulation, as mutations in the C-terminus prevented cell surface expression and abolished effects on channel gating. It is noteworthy that the location of the EGFP tag also profoundly affected the outcome of these experiments: tagging the C-terminal end of LMAN2 prevented cell surface localization and gating effects, whereas gating effects were observed when Kv1.2 was co-expressed with untagged LMAN2 or N-terminally tagged EGFP-LMAN2 (Fig. 8).

In terms of the relationship between redox and LMAN2 modulation of Kv1.2, it remains unclear whether LMAN2 influences Kv1.2 gating via a direct interaction, versus indirect effects on other regulatory proteins. Some other reports have highlighted potential effects of the σ-1 receptor (OPRS1) on pulse-dependent changes in Kv1.2 gating kinetics, along with activity-dependent trafficking effects *in vivo*, however these gating effects are less pronounced than effects observed for LMAN2 (Kourrich *et al*., 2013; Abraham *et al*., 2019). Kv1.1 and Kv1.2 channels also co-localize with a number of proteins (LGI1, ADAM22, ADAM23) in the juxtaparanodal region of myelinated axons, and the axon initial segment (Schulte *et al*., 2006; Ogawa *et al*., 2010; Kwon *et al*., 2016; Hivert *et al*., 2019; Fukata *et al*., 2021). The gating effects of these proteins alone or in combination with LMAN2 are not known. Another possible explanation for the effects of LMAN2 that is consistent with our data thus far is that LMAN2 may be involved in the maturation of a distinct regulatory protein that influences Kv1.2. We have not been able to consistently show a direct physical interaction between LMAN2 and Kv1.2 by approaches such as co-immunoprecipitation. Thus, there may be additional proteins or lipids involved in the effects reported here that remain to be discovered, or a delicate interaction that is broken in solubilization conditions used in our experiments. Overall, although there are very powerful functional outcomes that arise from co-expression of LMAN2 with Kv1.2, we see it as a strong likelihood that additional undiscovered regulators are involved in modulation of Kv1.2 activation properties.

In conclusion, we have detected TsTX-sensitive currents in DRG neurons, which exhibit prominent redox sensitivity similar to Kv1.2 channels expressed in heterologous systems. We have identified LMAN2 as a strong regulator of this sensitivity, as co-expression of Kv1.2 and LMAN2 mimics the effects of extracellular reducing agents (i.e. promotes slow activation kinetics) and sensitizes channels to extracellular reducing agents. Additionally, sensitivity to LMAN2 and redox is controlled by an overlapping region (intracellular S2-S3 linker) in Kv1.2. Overall, this study establishes LMAN2 as a strong candidate regulator of activation properties of Kv1.2, although we suspect the involvement of additional unidentified regulatory elements.

---

**Supplementary Figure 1.**
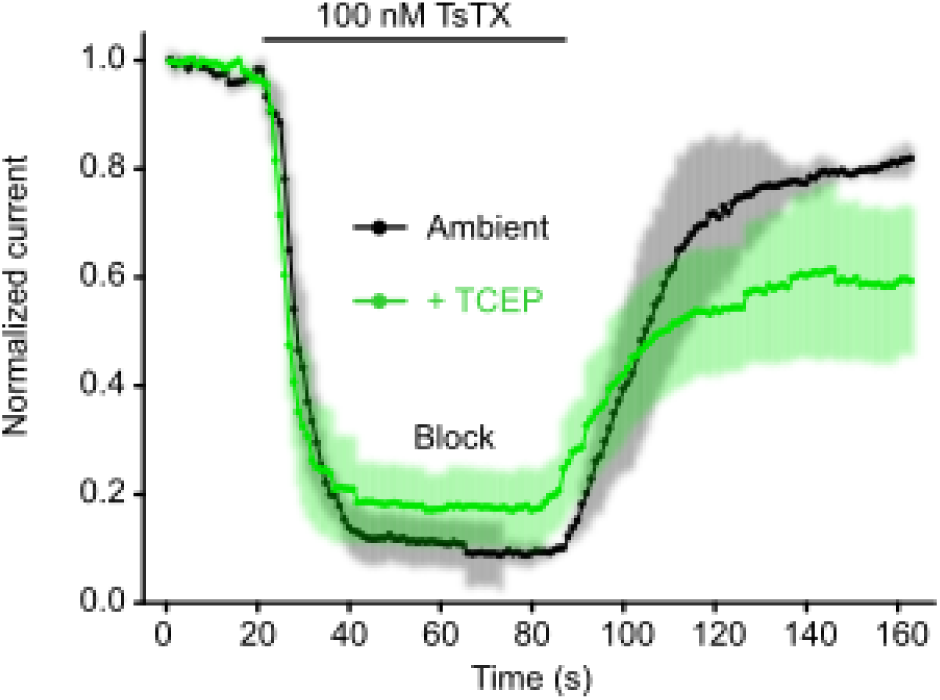
Tityustoxin-mediated Kv1.2 inhibition is retained in reducing conditions. Tityustoxin inhibition of Kv1.2 in a stable LM cell line was tested using patch clamp. Normalized current vs. time course of Kv1.2 channel block and washout with 100 nM TsTX from a stable Kv1.2 LM cell line in ambient redox (black, n = 4) or in the presence of 200 μM TCEP (green, n = 4). Depolarizing pulses to 30 mV (200 ms) were delivered at 1 Hz, in ambient redox conditions or 200 μM TCEP.

**Supplementary Figure 2.**
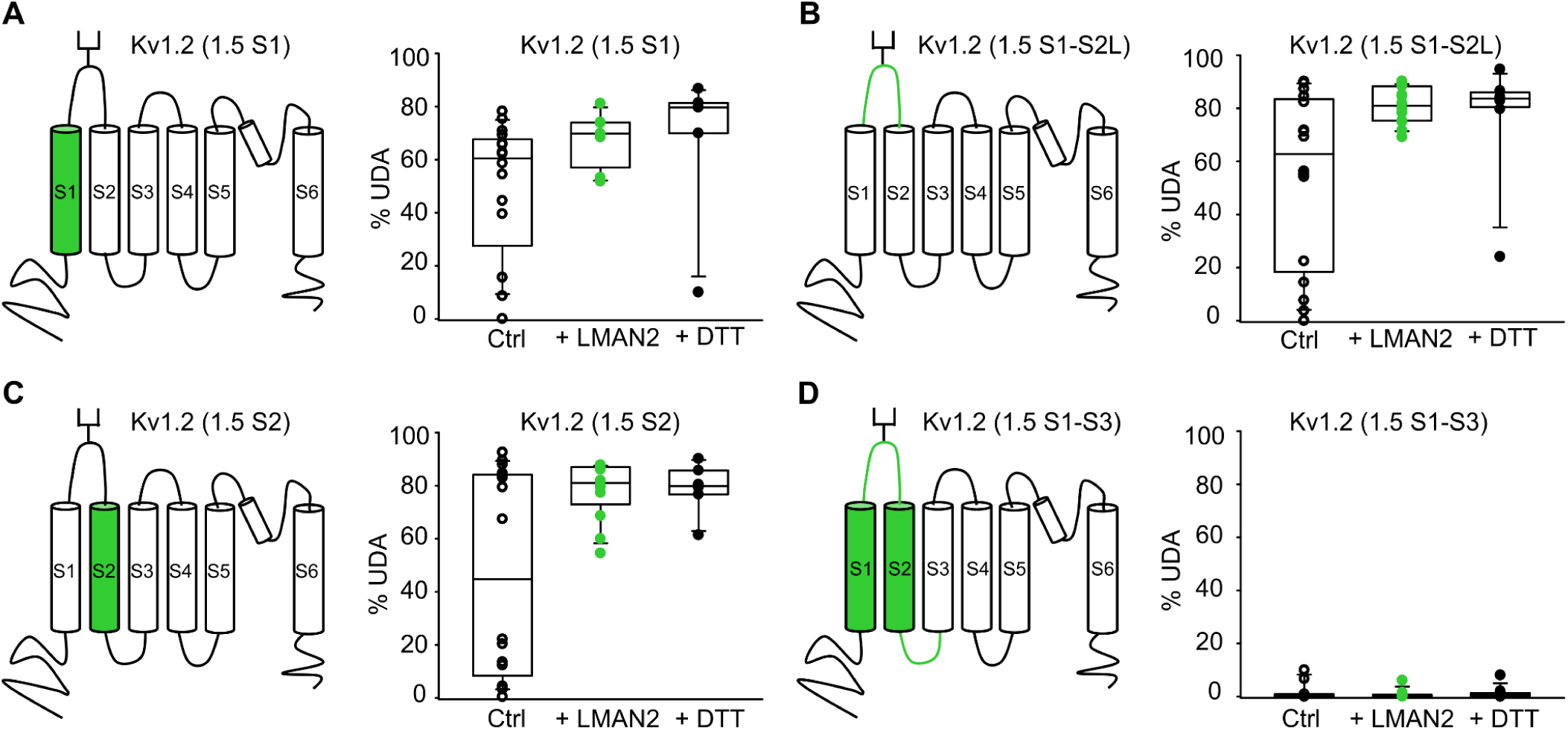
Chimeric analysis of Kv1.2 sensitivity to LMAN2 and DTT-mediated use-dependence activation. (A-D) Cartoons illustrating the chimeric channel design in which transmembrane or linker segments of Kv1.5 (green) were exchanged for the equivalent regions of Kv1.2 (black). (A-D) Summarized use-dependent activation (% UDA) of Kv1.2/1.5 chimeras in the presence or absence of LMAN2 or 200 μM DTT. Enhanced % UDA in the presence of LMAN2 and DTT was observed in all chimeras except the Kv1.2 (1.5 S1-S3) chimera. (Kv1.2 (1.5 S1), n = 6-16; Kv1.2 (1.5 S1-S2L), n = 7-16; Kv1.2 (1.5 S2), n = 6-16; Kv1.2 (1.5 S1-S3), n = 10).

## REFERENCES

Abraham MJ, Fleming KL, Raymond S, Wong AYC, and Bergeron R (2019) The sigma-1 receptor behaves as an atypical auxiliary subunit to modulate the functional characteristics of Kv1.2 channels expressed in HEK293 cells. Physiol Rep 7:e14147.

Arar C, Carpentier V, Le Caer JP, Monsigny M, Legrand A, and Roche AC (1995) ERGIC-53, a membrane protein of the endoplasmic reticulum-Golgi intermediate compartment, is identical to MR60, an intracellular mannose-specific lectin of myelomonocytic cells. J Biol Chem 270:3551–3553.

Baronas VA, McGuinness BR, Brigidi GS, Gomm Kolisko RN, Vilin YY, Kim RY, Lynn FC, Bamji SX, Yang R, and Kurata HT (2015) Use-Dependent Activation of Neuronal Kv1.2 Channel Complexes. J Neurosci 35:3515–3524.

Baronas VA, Yang R, Vilin YY, and Kurata HT (2016) Determinants of frequency-dependent regulation of Kv1.2-containing potassium channels. Channels Austin Tex 10:158–166.

Baronas VA, Yang RY, and Kurata HT (2017) Extracellular redox sensitivity of Kv1.2 potassium channels. Sci Rep 7:9142.

Baronas VA, Yang RY, Morales LC, Sipione S, and Kurata HT (2018) Slc7a5 regulates Kv1.2 channels and modifies functional outcomes of epilepsy-linked channel mutations. Nat Commun 9.

Brew HM, Gittelman JX, Silverstein RS, Hanks TD, Demas VP, Robinson LC, Robbins CA, McKee-Johnson J, Chiu SY, Messing A, and Tempel BL (2007) Seizures and reduced life span in mice lacking the potassium channel subunit Kv1.2, but hypoexcitability and enlarged Kv1 currents in auditory neurons. J Neurophysiol 98:1501–1525.

Buckley C, Oger J, Clover L, Tüzün E, Carpenter K, Jackson M, and Vincent A (2001) Potassium channel antibodies in two patients with reversible limbic encephalitis. Ann Neurol 50:73–78.

Corbett MA, Bellows ST, Li M, Carroll R, Micallef S, Carvill GL, Myers CT, Howell KB, Maljevic S, Lerche H, Gazina EV, Mefford HC, Bahlo M, Berkovic SF, Petrou S, Scheffer IE, and Gecz J (2016) Dominant KCNA2 mutation causes episodic ataxia and pharmacoresponsive epilepsy. Neurology 87:1975–1984.

Everill B, and Kocsis JD (1999) Reduction in potassium currents in identified cutaneous afferent dorsal root ganglion neurons after axotomy. J Neurophysiol 82:700–708.

Fan L, Guan X, Wang W, Zhao J-Y, Zhang H, Tiwari V, Hoffman PN, Li M, and Tao Y-X (2014) Impaired neuropathic pain and preserved acute pain in rats overexpressing voltage-gated potassium channel subunit Kv1.2 in primary afferent neurons. Mol Pain 10:8.

Fiedler K, Parton RG, Kellner R, Etzold T, and Simons K (1994) VIP36, a novel component of glycolipid rafts and exocytic carrier vesicles in epithelial cells. EMBO J 13:1729–1740.

Fiedler K, and Simons K (1994) A putative novel class of animal lectins in the secretory pathway homologous to leguminous lectins. Cell 77:625–626.

Fukata Y, Chen X, Chiken S, Hirano Y, Yamagata A, Inahashi H, Sanbo M, Sano H, Goto T, Hirabayashi M, Kornau H-C, Prüss H, Nambu A, Fukai S, Nicoll RA, and Fukata M (2021) LGI1-ADAM22-MAGUK configures transsynaptic nanoalignment for synaptic transmission and epilepsy prevention. Proc Natl Acad Sci U S A 118:e2022580118.

Füllekrug J, Scheiffele P, and Simons K (1999) VIP36 localisation to the early secretory pathway. J Cell Sci 112 ( Pt 17):2813–2821.

Goldberg EM, Clark BD, Zagha E, Nahmani M, Erisir A, and Rudy B (2008) K+ channels at the axon initial segment dampen near-threshold excitability of neocortical fast-spiking GABAergic interneurons. Neuron 58:387–400.

Grissmer S, Nguyen AN, Aiyar J, Hanson DC, Mather RJ, Gutman GA, Karmilowicz MJ, Auperin DD, and Chandy KG (1994) Pharmacological characterization of five cloned voltage-gated K+ channels, types Kv1.1, 1.2, 1.3, 1.5, and 3.1, stably expressed in mammalian cell lines. Mol Pharmacol 45:1227–1234.

Hara-Kuge S, Ohkura T, Ideo H, Shimada O, Atsumi S, and Yamashita K (2002) Involvement of VIP36 in intracellular transport and secretion of glycoproteins in polarized Madin-Darby canine kidney (MDCK) cells. J Biol Chem 277:16332–16339.

Hara-Kuge S, Seko A, Shimada O, Tosaka-Shimada H, and Yamashita K (2004) The binding of VIP36 and alpha-amylase in the secretory vesicles via high-mannose type glycans. Glycobiology 14:739–744.

Hauri H, Appenzeller C, Kuhn F, and Nufer O (2000) Lectins and traffic in the secretory pathway. FEBS Lett 476:32–37.

Hivert B, Marien L, Agbam KN, and Faivre-Sarrailh C (2019) ADAM22 and ADAM23 modulate the targeting of the Kv1 channel-associated protein LGI1 to the axon initial segment. J Cell Sci 132:jcs219774.

Hua Z, Watanabe R, Fukunaga T, Matsui Y, Matsuoka M, Yamaguchi S, Tanabe S-Y, Yamamoto M, Tamura-Kawakami K, Takagi J, Kajita M, Futai E, and Shirakabe K (2024) C-terminal amino acids in the type I transmembrane domain of L-type lectin VIP36 affect γ-secretase susceptibility. Biochem Biophys Res Commun 696:149504.

Ishikawa K, Tanaka M, Black JA, and Waxman SG (1999) Changes in expression of voltage-gated potassium channels in dorsal root ganglion neurons following axotomy. Muscle Nerve 22:502–507.

Itin C, Kappeler F, Linstedt AD, and Hauri HP (1995) A novel endocytosis signal related to the KKXX ER-retrieval signal. EMBO J 14:2250–2256.

Kamiya Y, Kamiya D, Yamamoto K, Nyfeler B, Hauri H-P, and Kato K (2008) Molecular Basis of Sugar Recognition by the Human L-type Lectins ERGIC-53, VIPL, and VIP36*. J Biol Chem 283:1857–1861.

Kamiya Y, Yamaguchi Y, Takahashi N, Arata Y, Kasai K-I, Ihara Y, Matsuo I, Ito Y, Yamamoto K, and Kato K (2005) Sugar-binding properties of VIP36, an intracellular animal lectin operating as a cargo receptor. J Biol Chem 280:37178–37182.

Kim DS, Choi JO, Rim HD, and Cho HJ (2002) Downregulation of voltage-gated potassium channel alpha gene expression in dorsal root ganglia following chronic constriction injury of the rat sciatic nerve. Brain Res Mol Brain Res 105:146–152.

Kourrich S, Hayashi T, Chuang J-Y, Tsai S-Y, Su T-P, and Bonci A (2013) Dynamic interaction between sigma-1 receptor and Kv1.2 shapes neuronal and behavioral responses to cocaine. Cell 152:236–247.

Kwon S-H, Oh S, Nacke M, Mostov KE, and Lipschutz JH (2016) Adaptor Protein CD2AP and L-type Lectin LMAN2 Regulate Exosome Cargo Protein Trafficking through the Golgi Complex. J Biol Chem 291:25462–25475.

Lamothe SM, and Kurata HT (2020) Slc7a5 alters Kvβ-mediated regulation of Kv1.2. J Gen Physiol 152:e201912524.

Lamothe SM, Sharmin N, Silver G, Satou M, Hao Y, Tateno T, Baronas VA, and Kurata HT (2020) Control of Slc7a5 sensitivity by the voltage-sensing domain of Kv1 channels. eLife 9:e54916.

Laumet G, Garriga J, Chen S-R, Zhang Y, Li D-P, Smith TM, Dong Y, Jelinek J, Cesaroni M, Issa J-P, and Pan H-L (2015) G9a is essential for epigenetic silencing of K(+) channel genes in acute-to-chronic pain transition. Nat Neurosci 18:1746–1755.

Lee S-M, Kim J-E, Sohn J-H, Choi H-C, Lee J-S, Kim S-H, Kim M-J, Choi I-G, and Kang T-C (2009) Down-regulation of delayed rectifier K+ channels in the hippocampus of seizure sensitive gerbils. Brain Res Bull 80:433–442.

Liang L, Gu X, Zhao J-Y, Wu S, Miao X, Xiao J, Mo K, Zhang J, Lutz BM, Bekker A, and Tao Y-X (2016) G9a participates in nerve injury-induced Kcna2 downregulation in primary sensory neurons. Sci Rep 6:37704, Nature Publishing Group.

Masnada S, Hedrich UBS, Gardella E, Schubert J, Kaiwar C, Klee EW, Lanpher BC, Gavrilova RH, Synofzik M, Bast T, Gorman K, King MD, Allen NM, Conroy J, Ben Zeev B, Tzadok M, Korff C, Dubois F, Ramsey K, Narayanan V, Serratosa JM, Giraldez BG, Helbig I, Marsh E, O’Brien M, Bergqvist CA, Binelli A, Porter B, Zaeyen E, Horovitz DD, Wolff M, Marjanovic D, Caglayan HS, Arslan M, Pena SDJ, Sisodiya SM, Balestrini S, Syrbe S, Veggiotti P, Lemke JR, Møller RS, Lerche H, and Rubboli G (2017) Clinical spectrum and genotype–phenotype associations of KCNA2-related encephalopathies. Brain 140:2337–2354, Oxford Academic.

McKinnon D (1989) Isolation of a cDNA clone coding for a putative second potassium channel indicates the existence of a gene family. J Biol Chem 264:8230–8236.

Muranaka M, Takamatsu S, Ouchida T, Kanazawa Y, Kondo J, Nakagawa T, Egashira Y, Fukagawa K, Gu J, Okamoto T, Kamada Y, and Miyoshi E (2023) Vesicular Integral-Membrane Protein 36 Is Involved in the Selective Secretion of Fucosylated Proteins into Bile Duct-like Structures in HepG2 Cells. Int J Mol Sci 24:7037.

Neve EPA, Svensson K, Fuxe J, and Pettersson RF (2003) VIPL, a VIP36-like membrane protein with a putative function in the export of glycoproteins from the endoplasmic reticulum☆. Exp Cell Res 288:70–83.

Nufer O, Mitrovic S, and Hauri H-P (2003) Profile-based data base scanning for animal L-type lectins and characterization of VIPL, a novel VIP36-like endoplasmic reticulum protein. J Biol Chem 278:15886–15896.

Ogawa Y, Oses-Prieto J, Kim MY, Horresh I, Peles E, Burlingame AL, Trimmer JS, Meijer D, and Rasband MN (2010) ADAM22, A Kv1 Channel-Interacting Protein, Recruits Membrane-Associated Guanylate Kinases to Juxtaparanodes of Myelinated Axons. J Neurosci 30:1038–1048.

Pena S d. j., and Coimbra R l. m. (2015) Ataxia and myoclonic epilepsy due to a heterozygous new mutation in KCNA2: proposal for a new channelopathy. Clin Genet 87:e1–e3.

Qin S-Y, Kawasaki N, Hu D, Tozawa H, Matsumoto N, and Yamamoto K (2012) Subcellular localization of ERGIC-53 under endoplasmic reticulum stress condition. Glycobiology 22:1709–1720.

Reiterer V, Nyfeler B, and Hauri H-P (2010) Role of the lectin VIP36 in post-ER quality control of human alpha1-antitrypsin. Traffic Cph Den 11:1044–1055.

Rettig J, Heinemann SH, Wunder F, Lorra C, Parcej DN, Oliver Dolly J, and Pongs O (1994) Inactivation properties of voltage-gated K+ channels altered by presence of β-subunit. Nature 369:289–294, Nature Publishing Group.

Rezazadeh S, Kurata HT, Claydon TW, Kehl SJ, and Fedida D (2007) An activation gating switch in Kv1.2 is localized to a threonine residue in the S2-S3 linker. Biophys J 93:4173–4186.

Sachdev M, Gaínza-Lein M, Tchapyjnikov D, Jiang Y-H, Loddenkemper T, and Mikati MA (2017) Novel clinical manifestations in patients with KCNA2 mutations. Seizure 51:74–76.

Satou M, Wang J, Nakano-Tateno T, Teramachi M, Suzuki T, Hayashi K, Lamothe S, Hao Y, Kurata H, Sugimoto H, Chik C, and Tateno T (2020) L-type amino acid transporter 1, LAT1, in growth hormone-producing pituitary tumor cells. Mol Cell Endocrinol 515:110868.

Schrag JD, Procopio DO, Cygler M, Thomas DY, and Bergeron JJM (2003) Lectin control of protein folding and sorting in the secretory pathway. Trends Biochem Sci 28:49–57.

Schulte U, Thumfart J-O, Klöcker N, Sailer CA, Bildl W, Biniossek M, Dehn D, Deller T, Eble S, Abbass K, Wangler T, Knaus H-G, and Fakler B (2006) The epilepsy-linked Lgi1 protein assembles into presynaptic Kv1 channels and inhibits inactivation by Kvbeta1. Neuron 49:697–706.

Shen W, Hernandez-Lopez S, Tkatch T, Held JE, and Surmeier DJ (2004) Kv1.2-containing K+ channels regulate subthreshold excitability of striatal medium spiny neurons. J Neurophysiol 91:1337–1349.

Sheng M, Liao YJ, Jan YN, and Jan LY (1993) Presynaptic A-current based on heteromultimeric K+ channels detected in vivo. Nature 365:72–75.

Shirakabe K, Hattori S, Seiki M, Koyasu S, and Okada Y (2011) VIP36 protein is a target of ectodomain shedding and regulates phagocytosis in macrophage Raw 264.7 cells. J Biol Chem 286:43154–43163.

Stirling L, Williams MR, and Morielli AD (2009) Dual Roles for RHOA/RHO-Kinase In the Regulated Trafficking of a Voltage-sensitive Potassium Channel. Mol Biol Cell 20:2991–3002.

Sun L, Gu X, Pan Z, Guo X, Liu J, Atianjoh FE, Wu S, Mo K, Xu B, Liang L, Bekker A, and Tao Y-X (2019) Contribution of DNMT1 to Neuropathic Pain Genesis Partially through Epigenetically Repressing Kcna2 in Primary Afferent Neurons. J Neurosci Off J Soc Neurosci 39:6595–6607.

Syrbe S, Hedrich UB, Riesch E, Djemie T, Muller S, Moller RS, Maher B, Hernandez-Hernandez L, Synofzik M, Caglayan HS, Arslan M, Serratosa JM, Nothnagel M, May P, Krause R, Loffler H, Detert K, Dorn T, Vogt H, Kramer G, Schols L, Mullis PE, Linnankivi T, Lehesjoki AE, Sterbova K, Craiu DC, Hoffman-Zacharska D, Korff CM, Weber YG, Steinlin M, Gallati S, Bertsche A, Bernhard MK, Merkenschlager A, Kiess W, Gonzalez M, Zuchner S, Palotie A, Suls A, de JP, Helbig I, Biskup S, Wolff M, Maljevic S, Schule R, Sisodiya SM, Weckhuysen S, Lerche H, and Lemke JR (2015) De novo loss- or gain-of-function mutations in KCNA2 cause epileptic encephalopathy. Nat Genet 47:393–399.

Wang H, Kunkel DD, Martin TM, Schwartzkroin PA, and Tempel BL (1993) Heteromultimeric K+ channels in terminal and juxtaparanodal regions of neurons. Nature 365:75–79.

Werkman TR, Gustafson TA, Rogowski RS, Blaustein MP, and Rogawski MA (1993) Tityustoxin-K alpha, a structurally novel and highly potent K+ channel peptide toxin, interacts with the alpha-dendrotoxin binding site on the cloned Kv1.2 K+ channel. Mol Pharmacol 44:430–436.

Xie G, Harrison J, Clapcote SJ, Huang Y, Zhang J-Y, Wang L-Y, and Roder JC (2010) A new Kv1.2 channelopathy underlying cerebellar ataxia. J Biol Chem 285:32160–32173.

Yamashita K, Hara-Kuge S, and Ohkura T (1999) Intracellular lectins associated with N-linked glycoprotein traffic. Biochim Biophys Acta 1473:147–160.

Yang E-K, Takimoto K, Hayashi Y, de Groat WC, and Yoshimura N (2004) Altered expression of potassium channel subunit mRNA and alpha-dendrotoxin sensitivity of potassium currents in rat dorsal root ganglion neurons after axotomy. Neuroscience 123:867–874.

Yang J-W, Vacher H, Park K-S, Clark E, and Trimmer JS (2007) Trafficking-dependent phosphorylation of Kv1.2 regulates voltage-gated potassium channel cell surface expression. Proc Natl Acad Sci 104:20055–20060, Proceedings of the National Academy of Sciences.

Zhang J, Rong L, Shao J, Zhang Y, Liu Y, Zhao S, Li L, Yu W, Zhang M, Ren X, Zhao Q, Zhu C, Luo H, Zang W, and Cao J (2021) Epigenetic restoration of voltage-gated potassium channel Kv1.2 alleviates nerve injury-induced neuropathic pain. J Neurochem 156:367–378.

Zhao J-Y, Liang L, Gu X, Li Z, Wu S, Sun L, Atianjoh FE, Feng J, Mo K, Jia S, Lutz BM, Bekker A, Nestler EJ, and Tao Y-X (2017) DNA methyltransferase DNMT3a contributes to neuropathic pain by repressing Kcna2 in primary afferent neurons. Nat Commun 8:14712, Nature Publishing Group.

Zhao X, Tang Z, Zhang H, Atianjoh FE, Zhao J-Y, Liang L, Wang W, Guan X, Kao S-C, Tiwari V, Gao Y-J, Hoffman PN, Cui H, Li M, Dong X, and Tao Y-X (2013) A long noncoding RNA contributes to neuropathic pain by silencing Kcna2 in primary afferent neurons. Nat Neurosci 16:1024–1031.

